# Condensin Accelerates Long-Range Intra-Chromosomal Interactions

**DOI:** 10.1101/2025.05.02.651983

**Authors:** Fan Zou, Yi Li, Timothy Földes, Henrik Dahl Pinholt, Courtney Smith, Leonid Mirny, Lu Bai

## Abstract

The 3D genome organization plays a key role in regulating interactions among chromosomal loci. While Chromosome Conformation Capture (3C)-based methods have provided static snapshots of chromatin architecture, the kinetics of chromosomal encounters in live cells remain poorly characterized. In this study, we employ Chemically Induced Chromosomal Interaction (CICI) to measure encounter times between multiple loci pairs in G1-arrested budding yeast. Our results show that chromosome motion closely follows the Rouse polymer model, with similar diffusion parameters at all tested loci. Surprisingly, we find that long-range intra-chromosomal encounters occur significantly faster than inter-chromosomal encounters at similar 3D distances. Using targeted depletion experiments, we identify condensin, but not cohesin, as the complex responsible for these rapid intra-chromosomal interactions. This is further supported by Hi-C analysis, which reveals that condensin promotes long-distance intra-chromosomal interactions in G1 yeast. Through polymer simulations, we estimate that condensin extrudes chromatin at ∼2 kb/s with a density of one complex per 1-2 Mb and a processivity of 120-220 kb. These findings uncover a novel role for condensin in shaping the interphase genome organization and provide new insights into chromosomal search dynamics *in vivo*.

## Introduction

Precise spatial and temporal regulation of nuclear processes, such as DNA replication, transcription, and repair, is essential for the survival of all organisms^1^. Increasing evidence highlights the critical role of 3D genome organization in orchestrating these processes^2^. Recent advances in Chromosomal Conformation Captures (3C)-based techniques and DNA Fluorescence In Situ Hybridization (FISH) assays have revealed intricate genome structures with unprecedented scale and resolution^3-5^. While highly insightful, these assays are typically conducted on fixed cells, providing only static snapshots of chromatin contacts rather than capturing the dynamics of chromosomal interactions in live cells.

The “encounter time” or “first passage time” between two loci, which reflects the speed at which different chromosomal regions interact with each other, is a key kinetic parameter for many genomic processes. For instance, the speed at which an enhancer locates its target promoter likely affects the rate of gene activation, while the time required for homologous search dictates the efficiency of double-strand break (DSB) repair. Given its functional significance, cells have evolved mechanisms to regulate chromosomal encounters. Chromosomes exhibit sub-diffusive motion in the nucleus^6^, and properties such as diffusion coefficient, confinement radius, and average 3D distance between the loci pair can have a major impact on their encounter rate. For example, DSBs increase the genome-wide chromosome mobility, which is correlated with more efficient homologous repair^7^. Another factor that can influence encounter rate is loop extrusion, a process driven by Structural Maintenance of Chromosomes (SMC) complexes, such as cohesin and condensin^8^. By extruding DNA until stalling at CTCF binding sites, cohesin is thought to promote chromosomal contacts within the topologically associating domains (TADs) while reducing contacts across TAD boundaries in higher eukaryotes^9^. Although SMC-mediated loop extrusions have been studied *in vitro*, mainly on naked DNA^10-12^, the speed, frequency, and processivity of loop extrusion in the chromatin context in live cells remain elusive. It is therefore unclear how loop extrusion quantitatively affects chromosomal interaction dynamics *in vivo*.

Accurate measurement of encounter times between chromosomal loci pairs is critical for understanding how cells regulate the interaction dynamics. In principle, this information can be obtained by fluorescently tagging two loci, tracking their motion through time-lapse imaging, and evaluating the time needed for the pair to approach within a certain proximity threshold. However, due to limited time and spatial resolution, this method may fail to capture every encounter, and the loci within the threshold may not form genuine interactions. A method we previously developed, Chemically Induced Chromosomal Interaction (CICI)^13^, allows us to overcome these limitations. By inserting LacO and TetO arrays into target genomic loci and expressing LacI-FKBP12 and TetR-FRB fusion proteins, stable interactions can be induced between the two loci upon rapamycin addition through FKBP12-FRB dimerization. The prolonged co-localization of the loci pair over time facilitates robust detection of encounter events and enables accurate quantification of encounter times.

In addition to experimental approaches, polymer physics provides valuable tools for studying chromosomal motion. In particular, the Rouse model, in which chromatin is approximated as a homogeneous series of beads connected by springs, has been shown to capture the main features of yeast chromosome dynamics^14-16^. Fitting Rouse model predictions to experimental data reveals chromatin properties like stiffness and persistence length. Incorporating loop extrusion into this framework and comparing it with experimentally measured encounter times allow us to infer key parameters of loop extrusion, including extrusion rate, frequency, and processivity.

In this study, we tracked the motion of 13 pairs of chromosomal loci in G1-arrested budding yeast through live-cell imaging and measured their encounter times using CICI, which we refer to as the CICI formation time (CFT). We found that the loci motion can be well-described by the Rouse polymer model, with diffusion parameters being largely homogeneous across all tested loci. Interestingly, CFTs of intra-chromosomal pairs are significantly shorter than those of inter-chromosomal pairs when separated by the same 3D distances, and only the inter-chromosomal encounter rates are consistent with the Rouse model prediction. Through targeted depletion experiments, we identified condensin, but not cohesin, as the key determinant of accelerated intra-chromosomal encounters in G1 yeast. Hi-C analysis further reveals that condensin depletion reduces intra-chromosomal contacts for regions separated by over ∼100 kb, supporting its role in promoting long-range intra-chromosomal interactions. Finally, by integrating the imaging and Hi-C data with loop extrusion simulations, we estimated that condensin extrudes chromatin with rates ∼2 kb/s, processivity between 120-200 kb, and a very low density of 1 condensin per 1-2 Mb. Overall, our findings identify a novel role for condensin in shaping interphase yeast genome organization and provide quantitative insights into how chromatin encounters are modulated inside cells.

## Results

### Measuring chromosome dynamics and encounter time of chromosomal loci pair using CICI

We previously developed CICI to induce interactions between two selected chromosomal loci^13^. In a rapamycin-resistant yeast strain, we express four fusion proteins, LacI-FKBP12, TetR-FRB, LacI-GFP, and TetR-mCherry, which bind to LacO and TetO arrays inserted into the target loci (**Figure 1A**). FKBP12 and FRB dimerize in the presence of rapamycin, leading to strong multivalent interactions between the two arrays. The GFP and mCherry fusion proteins enable visualization of the two arrays as green and red “chromatin dots”. By conducting time-lapse imaging of these dots in live cells, we can track the movement of the two loci and measure the CICI formation time (CFT). As chromatin dynamics and CICI formation are cell-cycle dependent^13,17^, all measurements below are carried out in G1-arrested cells (**Materials and Methods**). This also ensures that both LacO and TetO arrays are present as a single copy in each nucleus, simplifying subsequent data analysis.

**Figure 1.**
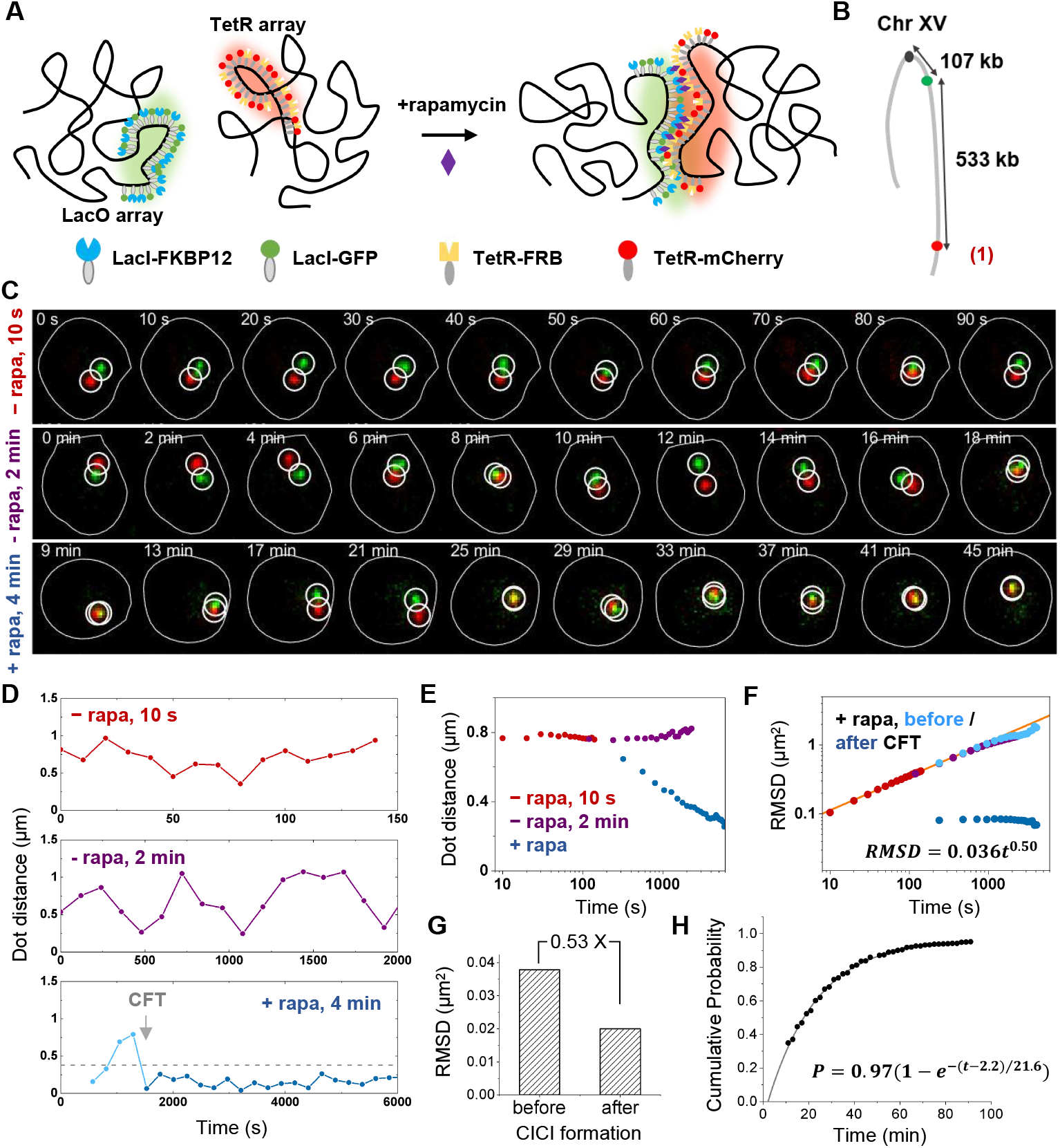
Measuring chromosome dynamics and encounter time of chromosomal loci pair using Chemically Induced Chromosomal Interaction (CICI). **A&B)** CICI scheme. The CICI strain contains LacO and TetO arrays (green and red dots in B) inserted into ChrXV at indicated locations (pair 1). Black dot represents the ChrXV centromere. **C)** Typical time-lapse images taken under three conditions: -rapamycin (-rapa) with 10 s interval (top row), -rapa with 2 min interval (middle), and +rapa with 4 min interval (bottom). Cell contours and detected dots are marked by white circles. **D)** Quantification of the dot distances in (C). CICI formation time (CFT) of the +rapa trace is marked by the gray arrow. After CFT, the two dots show continuous co-localization (distance below 0.38 µm marked by the dotted line). **E)** Population-averaged dot distances ±rapa. Red and purple: -rapa with two acquisition rates. Blue: +rapa. **F)** Relative mean square distance (RMSD) of the two dots ±rapa. Color scheme is the same as E except the +rapa data are split at CFTs (before: light blue, and after: dark blue). Power-law fit of the -rapa data is shown in the plot. **G)** Diffusion rate before and after CFT. The bar shows the mean RMSD at Δ*t* = 18 s. Note a 2-fold decrease of the RMSD after CICI formation. **H)** Cumulative probability of CFT over time. The dots are measurements, and the curve is the exponential fit.

We inserted the LacO and TetO into the right arm of ChrXV (separated by 533 kb) (pair 1) and performed time-lapse imaging with and without rapamycin at intervals of 4 and 2 minutes per frame, respectively (**Figure 1B & C**). To capture chromosome dynamics with higher temporal resolution, images were also acquired at 10-second intervals in the -rapamycin condition (**Figure 1C & D**). Distances between the LacO and TetO dots were extracted from these images (**Materials and Methods**). Without rapamycin, the two dots diffuse in the nucleus with fluctuating distances. Upon rapamycin addition, the dots initially display the same random motion before showing continuous co-localization, indicating CICI formation (**Figure 1D**). CICI formation is also supported by the population-averaged dot distance (**Figure 1E**). In the absence of rapamycin, the average distance remains ∼0.77 μm over time; following rapamycin addition, this distance progressively decreases as the two arrays become co-localized in a larger fraction of cells.

To quantify the diffusion properties of the arrays, we computed the relative mean squared displacement (RMSD) of the two dots as a function of time (**Materials and Methods**). The RMSD from the - rapamycin condition shows consistent time-dependence for images acquired at different frame rates (**Figure 1F**). These data at short time points are best fitted by a power law function, *RMSD(t) = 0*.*036t*^0.50^, with the exponent 0.5 perfectly matching the prediction of the Rouse polymer model^15^. In the long-time limit, RMSD is slightly lower than the *t*^1 / 2^ curve (**Figure 1F**), which is most likely due to the finite tethering by the DNA between the two loci and the confinement by the nuclear envelope. These results agree with previous MSD measurements of chromosome dynamics in yeast, showing that its chromosomes behave like a Rouse polymer^14-16,18^.

For the +rapamycin traces, we extracted the CFTs based on continuous co-localization and analyzed the traces before and after CFTs separately (**Materials and Methods**). The average RMSD before CFT is the same as that in the -rapamycin condition (**Figure 1F**), indicating that rapamycin does not affect the diffusion properties before the two arrays encounter. After CFT, RMSD becomes constant over time as the relative distance between the two dots is constrained (**Figure 1F**). Taking advantage of the short lag time between the acquisition of red and green images, we can use the RMSD value to estimate the diffusion constant after CICI formation (**Materials and Methods**). Remarkably, the diffusion rate of the interacting arrays is ∼50% lower than the unlinked ones (**Figure 1G**), which is exactly what we expect based on the Rouse model and Langevin equation (**Materials and Methods**).

The time-lapse imaging cannot capture CICI formations that occur too fast (<10 min) or too slow (>95 min). For example, out of the 1026 cells that reliably yield CFTs, 257 have already formed CICI in the first frame (7-10 min post rapamycin addition), and 50 show no CICI formation till the end of the movie. To quantify CFT more accurately, we plotted the cumulative CFT probability of the cell population (**Figure 1H**). This histogram can be well-fitted by a single exponential function *y* = 0.97(1 - *e*^-(*t*-2.2)/21.6^), indicating that the average CFT for this loci pair is 21.6 ± 0.4 min. Overall, these data demonstrate that the CICI method can be used to measure chromosome dynamics and encounter times between selected loci.

### CFTs between intra- and inter-chromosomal pairs have distinct distance-dependence

To understand how CFTs change with variables such as loci distance and diffusion constant, we extended our analysis in Figure 1 to 11 other loci pairs (pair 2-12) on ChrIV or ChrXV (**Figure 2A**). These loci pairs were selected to cover different configurations (intra-chromosomal on the same or different arms, or inter-chromosomal), linear distances, and nucleus localizations (based on the Rabl configuration). We imaged these loci in G1 cells ± rapamycin to measure their diffusion and CICI formation kinetics. The RMSDs of all these pairs exhibit a time-dependence similar to the pair measured in **Figure 1**, with exponents around 0.5 and scaling factors between 0.039 - 0.053 μm^2^/s (**Figure 2B & C, S1A**). We therefore conclude that diffusion dynamics are relatively homogeneous across all tested loci.

**Figure 2.**
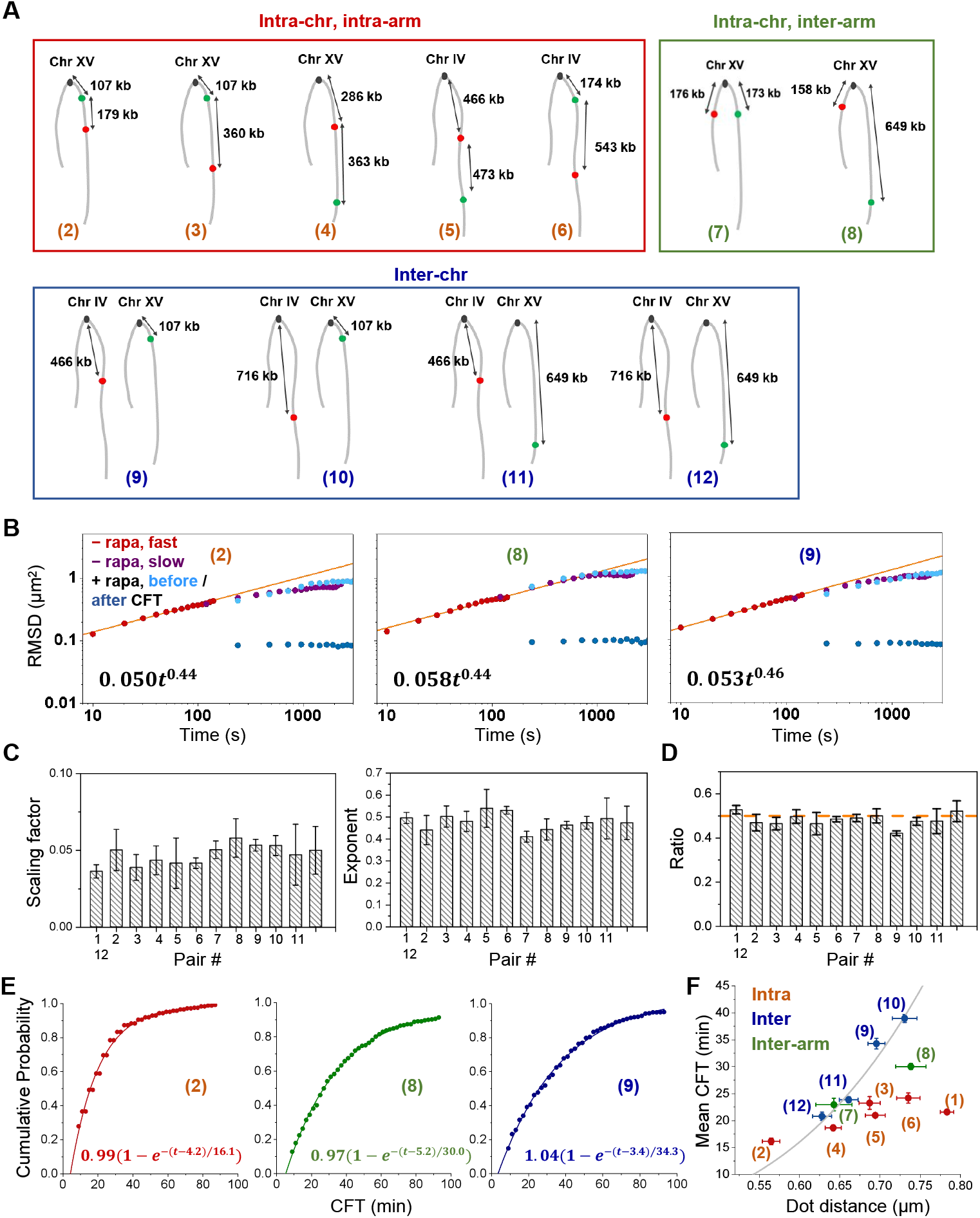
CFTs between intra- and inter-chromosomal pairs have distinct distance-dependence. **A)** Configurations of CICI pairs 2-12, which are divided into intra-arm, inter-arm, and inter-chromosomal groups. **B)** RMSDs as a function of time for pair 2, 8, and 9. Other pairs are shown in Figure S1A. The color scheme is the same as in Figure 1F. **C)** The best-fit scaling factor and exponent of the RMSDs for all pairs. Y error bars represent 95% confidence interval. **D)** The ratio of the RMSDs before and after CICI formation (after / before) for all pairs. Orange line marks the 0.5 ratio. **E)** Cumulative CFT probabilities for pair 2, 8, and 9. The single exponential fit is shown in each panel. Other pairs are shown in Figure S1B. **F)** Mean CFTs as a function of mean dot distances. The grey curve is the fit of the inter-chromosomal data with the power law: *CFT* ∝(*dis* −0.1*μm*)^3.8^. Y error bars represent 95% confidence interval of the CFT fit, and the X error bars are the standard error of the dot distance.

After rapamycin addition, CICI is successfully induced in a large fraction of cells within 90 min across all tested loci pairs. The diffusion rates of the co-localized LacO / TetO dots drop to ∼50% in comparison to the unpaired ones, further supporting the formation of CICI (**Figure 2D**). The cumulative CFT curves for all pairs follow single exponential functions but with distinct time constants (**Figure 2E, S1B**). Since diffusion properties of all tested loci are largely uniform, the variable CICI formation times should primarily reflect variations in the 3D distances between loci pairs. To illustrate this, we plotted the CFTs against the average distance between each pair (**Figure 2F**). Interestingly, intra-arm and inter-chromosomal CFTs display markedly different distance dependencies: loci within the same chromosome arm encounter each other significantly faster than loci on different chromosomes, even when their average physical separations are comparable. The inter-arm pairs (pairs 7 and 8) exhibit intermediate CFTs between those of intra-arm and inter-chromosomal pairs (**Figure 2F**).

For two freely diffusing dots with *MSD* ∝ *t*^1/2^, the average encounter time is expected to scale with the fourth power of the mean distance between the two dots, i.e. *CFT* ∝ *dis*^4^ (**Materials and Methods**). Due to confinement, the encounter time should be faster than in free solution as the region to explore is more restricted. CFTs of the inter-chromosomal pairs can be best fitted by *CFT* ∝ (*dis* - 0.1 μ*m*)^3.8^ (grey curve in **Figure 2F**), which agrees well with the theoretical prediction (0.1 μm is likely due to the finite size of the two arrays). CFTs between intra-chromosomal pairs, however, are much less sensitive to distance (**Figure 2F**). When separated by the same 3D distance over 0.65 µm, intra-chromosomal encounters occur significantly faster than the inter-chromosomal ones. Note that all the intra-chromosomal pairs tested here are over 150 kb and 0.5 μm apart, well above the previously estimated persistence length of the interphase yeast chromatin (10-20 kb and 0.05-0.2 μm)^19^. Therefore, despite the physical linkage, the two loci on the same chromosome should show move independently. If encounters were purely diffusion-mediated, we would expect intra- and inter-chromosomal pairs to behave similarly at comparable distances. The observed difference suggests that intra-chromosomal encounters are selectively facilitated by an additional mechanism.

### Condensin, but not cohesin, accelerates intra-chromosomal interactions in G1 cells

Loop extrusion by SMC complexes may account for the observation in **Figure 2F**. By translocating along DNA and forming loops within individual chromosomes, this activity could selectively promote the encounter of intra-chromosomal loci. It may also explain the reduced effect on inter-arm pairs, as centromeres are known to act as barriers to loop extrusion^20^. However, two major loop extruding complexes, cohesin and condensin, are both thought to have low activities in G1 yeast. Cohesin subunits are transcriptionally repressed in G1, and they fail to load efficiently onto chromosomes due to continuous cleavage by the Esp1 separin^21^. The level of functional condensin complexes is also reduced in G1 due to the limiting Cap-G subunit, YCG1^22^. Moreover, condensin distribution on the genome is largely restricted to specialized regions, such as rDNA, centromere, and recombination enhancer (RE) locus^23,24^. Therefore, it is not clear if either complex plays a role in CICI formation kinetics.

To test the function of SMC complexes in CICI formation kinetics, we rapidly depleted Smc1 or Smc4, the ATPase subunits of cohesin and condensin, respectively, and measured the resulting CFTs in G1 cells (**Materials and Methods**). Consistent with previous reports^25,26^, Smc1 tagged with auxin-inducible degron (AID) shows partial degradation even in the absence of auxin (IAA), with protein level reduced to ∼71% of WT (**Figure 3A**). After 1 hr of IAA treatment, Smc1 level further drops to ∼26% of WT. We compared CFTs across all three conditions: WT, AID -IAA, and AID +IAA. Smc1 depletion does not significantly affect the motion of pairs 1 and 4 in the absence of rapamycin (**Figure S2**); the CFTs and average distances of these pairs also remain similar across all conditions (**Figure 3B**). Notably, pair 1, which forms CICI much faster than inter-chromosomal pairs with similar 3D distances (**Figure 2F**), retains the same rate in the absence of cohesin. These data argue that cohesin is not responsible for the rapid CICI formation observed between intra-chromosomal loci.

**Figure 3.**
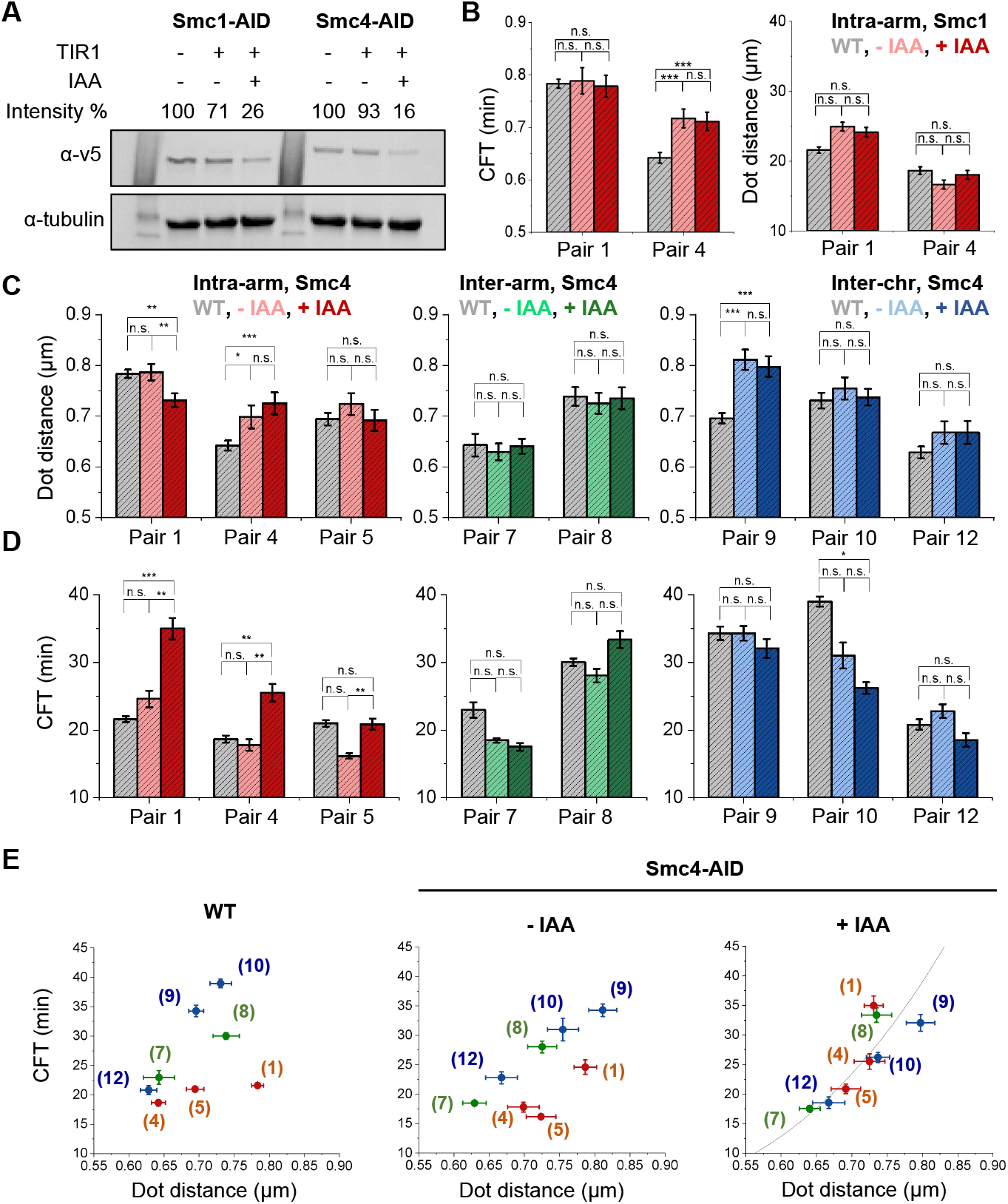
Condensin, but not cohesin, accelerates intra-chromosomal interactions in G1 cells. **A)** Western blot of V5-tagged Smc1 and Smc4 in a -TIR1 (E3 ligase) strain, or a +TIR1 strain with or without 1hr IAA treatment. The undigested percentage relative to the –TIR1 control is shown (average of two biological replicates). **B)** CFTs (left) and mean dot distances (right) for pair 1 and 4 in WT and Smc1-AID strain ±IAA. **C&D)** Mean dot distances (top) and CFTs (bottom) for the labeled pairs in WT and Smc4-AID strain ±IAA. **E)** CFT as a function of mean dot distance for labeled pairs in WT (left), Smc4-AID strain –IAA (mid), and Smc4-AID strain +IAA (right). Y error bars represent 95% confidence interval of the CFT fit, and the X error bars are the standard error of the dot distance. The grey curve is the fit of all data shown here with the power law: *CFT* ∝(*dis* −0.1*μm*)^3.3^. For CFT, statistical significance was determined using bootstrap resampling with replacement for 100000 iterations. For dot distance, statistical analysis is based on two-tailed students’ t-test, ***P<0.001, **P<0.01, *P<0.05, n.s. non-significant.

We next carried out similar measurements in the absence of condensin subunit Smc4. Smc4-AID is depleted to ∼93% of the WT level in -IAA and reaches ∼18% in +IAA (**Figure 3A**). We first evaluated the effect on pair 1, 4, and 5 with the two loci located on the same chromosome arm (**Figure 2A**). Depletion of condensin does not significantly affect the dynamics of these loci in the absence of rapamycin (**Figure S3**). The effect on the mean distances between these pairs is also small (**Figure 3C**). Strikingly, we observed large increases in CFTs after IAA treatment in comparison to the WT and - IAA conditions for pair 1 (from ∼22 to ∼35 min, *P*-value = 8.0e-5) and pair 4 (from ∼18 to ∼26 min, *P*-value = 2.7e-3) (**Figure 3D & S4**). For an unknown reason, CFT of the pair 5 is shorter in the Smc4-AID strain in the absence of IAA than in WT cells. Nevertheless, treatment with IAA in the AID strain still significantly increased CFT from ∼16 to ∼21 min (*P*-value = 9.8e-3). We also performed the same measurements on the inter-arm pairs 7 and 8. Pair 8, which has a longer 3D distance, shows a moderate and statistically insignificant increase upon condensin depletion (**Figure 3D & S4**). These data indicate that condensin accelerates intra-chromosomal encounters, especially for loci on the same chromosome arm.

To determine whether the condensin-dependent acceleration of contact formation is specific to intra-chromosomal interactions, we extended our analysis to inter-chromosomal pairs 9, 10, and 12. For all three pairs, we observed a mild, statistically insignificant decrease in CFTs in the +IAA condition compared to -IAA (**Figure 3D & S4**). In other words, the effect of condensin on inter-chromosomal encounters is small, and if anything, the trend is in the opposite direction of what we observed for intra-chromosomal pairs.

As a summary, we plotted CFT as a function of mean dot distance for WT and Smc4-AID strains with or without IAA for all the pairs tested (**Figure 3E**). The difference between intra- and inter-pairs becomes smaller in the Smc4-AID strains, even in the absence of IAA, likely due to the background degradation of Smc4. Strikingly, this difference is abolished in the +IAA condition with an overall fit of *CFT* ∝ (*dis* - 0.1 μ*m*)^3.3^ (**Figure 3E**). Together, these results support a model where all loci can rely on diffusion to establish interactions, but intra-chromosomal, especially intra-arm, communications can be further accelerated by loop extrusion.

### Condensin depletion reduces genome-wide long-distance intra-chromosomal contacts in G1 yeast

Most studies of condensin focus on M phase and/or at specific genomic loci. Even in mitotic yeast cells, depletion of condensin only has a mild effect on genome-wide contact frequencies^27^. Since our data support condensin-mediated loop extrusion during G1, condensin may also affect yeast interphase genome organization. To test this idea, we performed Hi-C experiments on WT and Smc4-AID strains in G1-arrested cells (**Figure S5A & B**). Hi-C signals show minor changes in the Smc4-AID strain compared to WT in the absence of IAA, likely reflecting the basal degradation of Smc4. More pronounced changes in Hi-C profiles is observed after IAA treatment (**Figure S5A & B**). We therefore focus on the comparison between the high degradation condition (Smc4-AID, +IAA) and WT in the following analyses.

When averaged across all chromosomes, condensin depletion results in decreased intra-chromosomal interactions (**Figure S5C**). For example, the average Hi-C signals for loci pairs separated by 200 kb (both intra-arm and inter-arm) are reduced by ∼15% in the absence of condensin (**Figure S5C**). This is consistent with our observation that condensin accelerates interactions between intra-chromosomal loci pairs. Notably, the impact of condensin depletion varies among individual chromosome arms. For the right arms of ChrIV and XV, where most CFT measurements were performed, condensin depletion leads to a greater reduction in Hi-C signals on ChrXV (16% over 200 kb) than ChrIV (12% over 200 kb) (**Figure 4A & B**). This difference may be due to variations in condensin density, as Smc4 and Brn1 (another condensin subunit) ChIP-exo data show higher enrichment of condensin binding over ChrXV than ChrIV (**Figure 4C**)^28^. This may also explain the smaller change of CFT for pair 5 after condensin depletion (**Figure 3E**).

**Figure 4.**
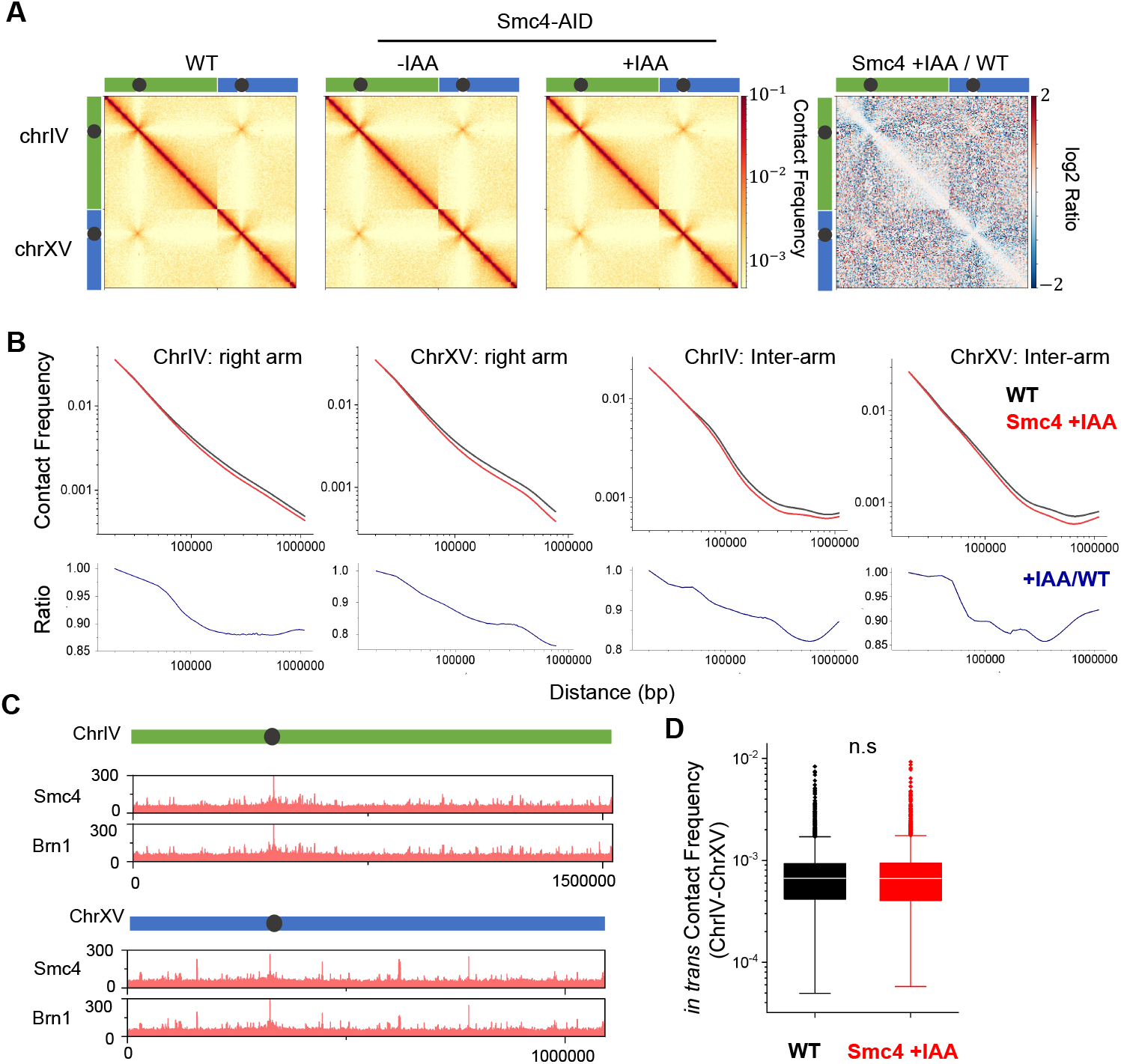
Condensin regulates the interphase budding yeast genome. **A)** Hi-C contact matrices over chrIV and chrXV in G1-arrested WT and Smc4-AID strain ±IAA. Log2 fold change between Smc4-AID +IAA and WT is shown on the right. **B)** Average pair-wise contact frequencies as a function of genomic distances in WT or Smc4-depleted conditions. Region pairs within the right arms of chrIV and ChrXV, or across between the left and right arms, are used for this calculation. The lower panels show the Smc4 +IAA / WT ratio. **C)** ChIP-exo data of Smc4 and Brn1 (another condensin subunit) on ChrIV and ChrXV. **D)** Inter-chromosomal contact frequencies between ChrIV and ChrXV in G1-arrested WT strain and Smc4-AID strain +IAA.

In contrast to intra-chromosomal interactions, inter-chromosomal interactions remain largely unaffected by condensin depletion except for a few specialized regions (see below). There is no significant difference in inter-chromosomal contact frequencies when pooled over all chromosomes or between ChrXV and ChrIV (**Figure 4D & S5D**). This is again consistent with the small and non-significant effect of condensin depletion on inter-chromosomal CFTs observed above (**Figure 3D**).

Consistent with previous findings that condensin is enriched over centromeres, RE, and rDNA, we found that condensin depletion has particularly pronounced effects on the conformation of pericentric regions, ChrIII, and ChrXII, respectively (**Figure S5A & B**). Within ∼200 kb of centromeres, condensin depletion reduces the interactions between centromeres and the flanking regions *in cis* but enhances the interactions between different centromeres *in trans* (**Figure S6A**). This pattern supports a model where condensin compacts and individualizes pericentromeric regions. Similar effects have been reported in mitotically arrested cells^27^, and our results suggest this centromeric organization is maintained during G1. On ChrIII, which contains all loci involved in the mating-type switch, condensin depletion led to a broad reduction in interactions between the left and right arms, including the key contacts between RE, *HML*, and the *MAT* locus (**Figure 5A**). This observation aligns with condensin’s known function in promoting mating-type switch between *MAT* and *HML*^24,29,30^.

**Figure 5.**
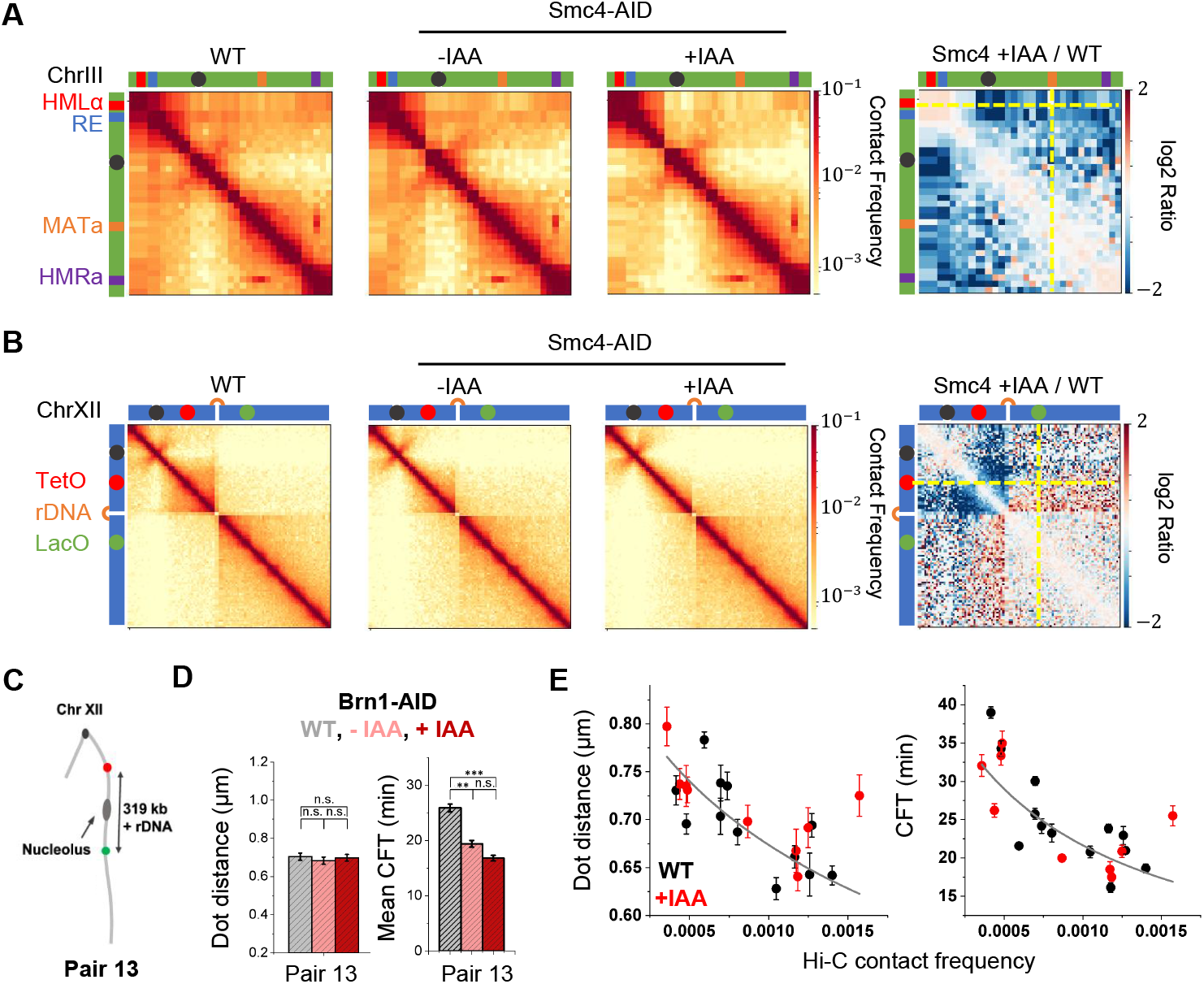
Condensin has pronounced effect on the conformation of ChrIII and ChrXII. **A&B)** Hi-C contact matrices for ChrIII (A) and ChrXII (B) in G1-arrested WT strain and Smc4-AID strain ±IAA. Key loci on these chromosomes, including centromeres (block dot), *HML*, RE, *MAT, HMR*, and rDNA, are labeled. Log2 fold change between Smc4-AID +IAA and WT is shown on the right. The intersection of the two yellow dashed lines in A marks the interaction between *HML*/RE and *MAT*, which is decreased in Smc4 +IAA. The one in B corresponds to the interaction between CICI pair 13 (C), which is increased in Smc4 +IAA. **C)** Schematic configuration of intra-chromosomal CICI pair 13, with one array on each side of rDNA. **D)** Mean dot distance and CFT for pair 13 in WT and Brn1-AID strain ±IAA. Brn1 (another condensin subunit) is depleted instead of Smc4 because the *SMC4* gene is located very close to the LacO array of pair 13, making it difficult to genetically manipulate without interfering with the locus. Same statistical test is used as in Figure 3. **E)** Mean dot distance and CFT vs Hi-C contact frequency for all CICI pairs tested. The data taken in WT and Smc4+IAA condition are shown in black and red. The power-law fits of these data are shown in grey curves: *dis* = 0.068(*cf* + 0.0008)^−0.33^ + 0.1, where 0.1 μm represents the finite size of the arrays (same as in Figure 2F), and *CFT* = 0.075(*cf* + 0.0008)^−0.9^.

On ChrXII, which harbors the rDNA repeats, condensin depletion significantly reduces the self-association of a region between the centromere and rDNA repeats, allowing it to interact more with the rest of ChrXII *in cis* and other chromosomes *in trans* (**Figure 5B & S5B**). This effect is already evident in the -IAA condition and is only moderately enhanced with IAA treatment (**Figure 5B & S5B**). These results suggest that the region between the centromere and rDNA is normally condensed and insulated from the rest of the genome^27,31^. This insulated state seems to be very sensitive to condensin concentration and is disrupted even with mild condensin depletion in -IAA. Supporting this model, an intra-chromosomal CICI pair spanning the rDNA locus (pair 13) exhibits an accelerated encounter rate in both -IAA and +IAA conditions (**Figure 5C & D**), and the majority of this acceleration already occurs in -IAA. Notably, this effect is in stark contrast to the trend observed for other intra-chromosomal pairs, where condensin depletion typically slows encounter kinetics (**Figure 3E**).

To further connect our imaging and Hi-C measurements, we plotted the CFTs and mean dot distances (*dis*) against the corresponding Hi-C contact frequencies (*cf*) for all CICI pairs tested. As expected, higher contact frequencies are associated with shorter dot distances and faster encounters (**Figure 5E**). These trends hold true in both WT and condensin-depleted cells. Quantitatively, CFT and 3D distance should follow power-law relationships with contact frequencies: *CFT* (*dis*) ∝ *cf*^*α*(*β*)^, where the *CFT* exponent should be around -1, and the *dis* exponent *β* should be between -0.25 and -0.3^32^. After adjusting *cf* by adding a constant to correct for the baseline level, we found that our data yield best-fit parameters of *α* ≈ -0.9 and *β* ≈ 0.33 (**Figure 5E**), in excellent agreement with theoretical predictions.

### Polymer simulations with sparse extruders recapitulate enhanced search time and moderate compaction

Our experiments show that condensin depletion significantly prolongs search times between distal intra-chromosomal loci, while having only a modest effect on Hi-C contact frequencies and 3D distances. To understand how condensin can accelerate search, we developed a polymer model of yeast chromosomes with active loop extrusion with the following goals: 1) to determine whether loop extrusion by condensin can accelerate search, while having little effect on average separations and Hi-C, 2) to estimate the parameters of condensin-mediated loop extrusion in yeast, and 3) to gain insights into the role of loop extrusion in facilitating long-range genomic interactions with implications for other organisms, e.g. enhancer-promoter communication in vertebrates.

Largely unconstrained polymers were shown to be adequate models for yeast chromosome organization^15,33^ and dynamics^14,16^. We confirmed this in our system by showing that experimentally measured MSD values for pairs of loci separated by different genomic distances are perfectly fit by the theoretical two-point MSD for the Rouse chain (**Figure S7A**). Thus, we modeled chromosomes as identical, flexible bead-and-spring polymers diffusing within a spherical volume with 1.5 µm radius at a physiological volume density of 15%. For WT simulation, we also incorporated condensin-mediated loop extrusion (see below).

We first calibrated our model *without loop extrusion* to reproduce experimentally measured quantities: diffusion dynamics, average distances, and the search times in condensin-depleted cells (+IAA experiments) (see **Methods** and **Figure S7B & S7C**). Once calibrated, we introduced loop extrusion characterized by the velocity of extrusion (*ν*), the average loop size (*λ*), and the average separation of extruders (*d*). We then assessed how extrusion affected search times, mean distances, and contact probabilities between loci pairs, comparing the results to experiments with condensin (-IAA).

The modest effect that we observe of condensin depletion on Hi-C contact frequencies led us to explore the sparse loop extrusion regime, where the separation between extruders is much larger than the average loop size (*d*>>*λ*, **Figure 6A & B**)^34^. In this regime, only a small fraction of the chain is extruded into loops, leaving large gaps of free polymer in between. We explored extrusion velocities in the range consistent with *in vitro*^10^ and *in vivo* estimates^35^, 1-4 kb/s, and performed polymer simulations with variable loop sizes and separation at low loop density (*λ*/*d* between 0.05 and 0.2) (**Figure 6C-E, S8**). As expected, we found that increasing the coverage by extruded loop (*λ*/*d*) leads to smaller distances and faster search (different curves on **Figure 6C & D**). Importantly, the effect on the search time is much more pronounced (15-40% reduction) than the effect on pairwise distances (3-10% reduction) (**Figure 6E**). Consistent with experimental data, we also found that loop extrusion does not accelerate trans-chromosomal interactions (**Figure 6F & G**). We conclude that sparse loop extrusion could indeed underlie the observed effects.

**Figure 6.**
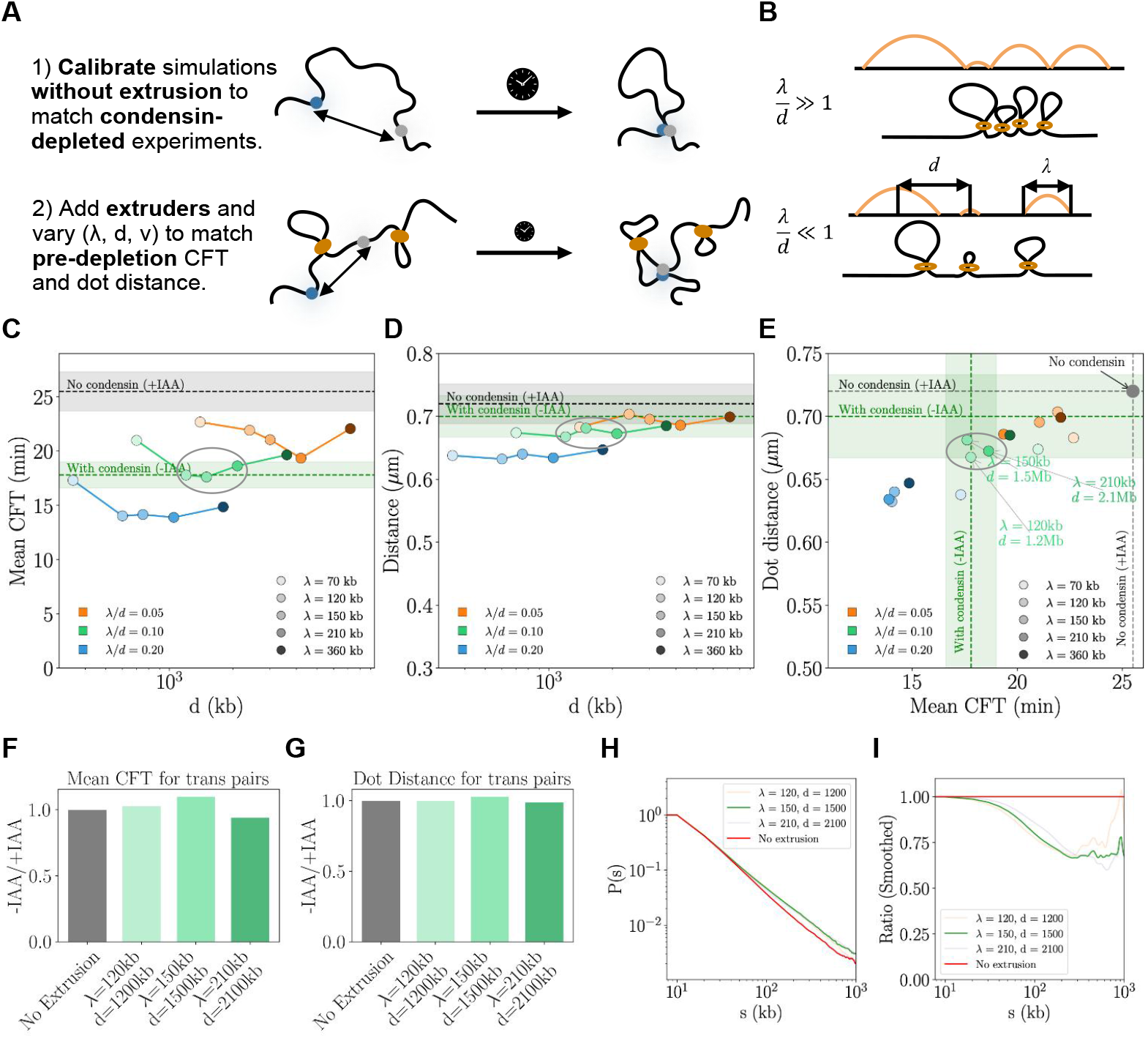
Polymer simulations with sparse extruders recapitulate enhanced search time and moderate compaction. **A)** Procedure of simulation. **B)** Schematic representation of chromatin chain looped by extruders, and the corresponding 1D arch diagram for a dense loop conformation (top) and sparse conformation (bottom). The average separation between loops *d*and loop size *λ* are depicted. **C & D)** Average search time (C) and distance (D) between two monomers separated by 360 kb (corresponding to the genomic separation of pair 4) with loop extrusion at *ν* = 2*kb* .*s*^−1^, shown for different values of loop size *λ* and loop separation *d*. Dashed lines indicate the experimentally measured search times in cells with condensin (-IAA, green) and without condensin (+IAA, grey), with shaded regions representing the corresponding error bars. **E)** The same simulation results as in C & D, but with dot distance plotted against search time. Simulations where both the dot distance and search time fall within 1.5 times the experimental error are annotated. **F & G)** CFT (F) and and average distance (G) for simulated *trans* pairs with best matching extrusion parameters. **H)** Simulated contact frequency as a function of genomic distance with extrusion that matched experimental data (orange, green, purple) and without extrusion (red). **I)** Ratio of simulated contact frequency (-extrusion divided by +extrusion) using the same parameters as in H.

Next, we sought to achieve quantitative agreement between the model and experimental data for pair 4, where condensin-mediated acceleration is most pronounced and the arrays are positioned far from the centromere. We found that reproducing experimental CFT and distance data together put considerable constraints on the loop extrusion parameters. The best agreement with experimental data was obtained with an average condensin loop size of *λ* ≈ 120-220 kb, and the average density of extruders of 1 per *d* ≈ 1-2 Mb, yet always with the loop coverage of *λ*/*d* ≈1/10 (**Figure 6E**, green shading). Interestingly, we also found that the velocity of extrusion *ν* should be relatively high, ≥ 2 kb/s, as simulations with slower velocities consistently failed to reproduce the experimental data (**Figure S8A-C**).

We also found that the best-fit parameters (*ν* = 2 kb/s and *λ/d* = 0.1) can reproduce changes in Hi-C contact probability upon condensin loss (for the arm containing pair 4), with a somewhat larger effect in simulations than in Hi-C (the fold change peaking at ∼35% vs. ∼20% in Hi-C). Thus we found that loops of yeast condensin are comparable to those of vertebrate cohesin, yet they are much sparse in yeast G1, as was suggested earlier by models that fit Hi-C data^27^. The processivity (*λ*) is also comparable to that of vertebrate condensin I, and speeds are similar to vertebrate condensins^35,36^.

Finally, our simulations provide mechanistic insight into *how* condensin-mediated loop extrusion accelerates long-distance contacts between chromosomal elements. We identify two classes of processes that enhance search efficiency (**Figure 7A**). The first involves a single extruder landing between the two elements and extruding a substantial portion of the intervening genome, thereby bringing the elements into close proximity. Such a large extrusion event allows the two elements to find each other rapidly (∼1-3 min) (**Figure 7B & S9A & S9C**). The second mechanism is due to multiple extrusion events that all moderately shorten the linker, collectively compacting the fiber everywhere and thus accelerating the search (**Figure 7C & S9A & S9C**). In that case, extrusion events are not immediately followed by contact formation (**Figure 7C**). While the first mechanism leads to rapid search, such large extrusion events are rare; small extrusion events are more frequent, but their individual contributions are small (**Figure 7D**). To dissect the effect of different extrusion size, we performed simulations where we selectively removed extrusion events of specific sizes (Methods & **Figure S9B**). These analyses reveal that both small and large events contribute to faster search, with large events playing a disproportionately important role (**Figure 7D**). Ultimately, it is the combined effect of both mechanisms that drives the observed acceleration in chromosomal encounter.

**Figure 7.**
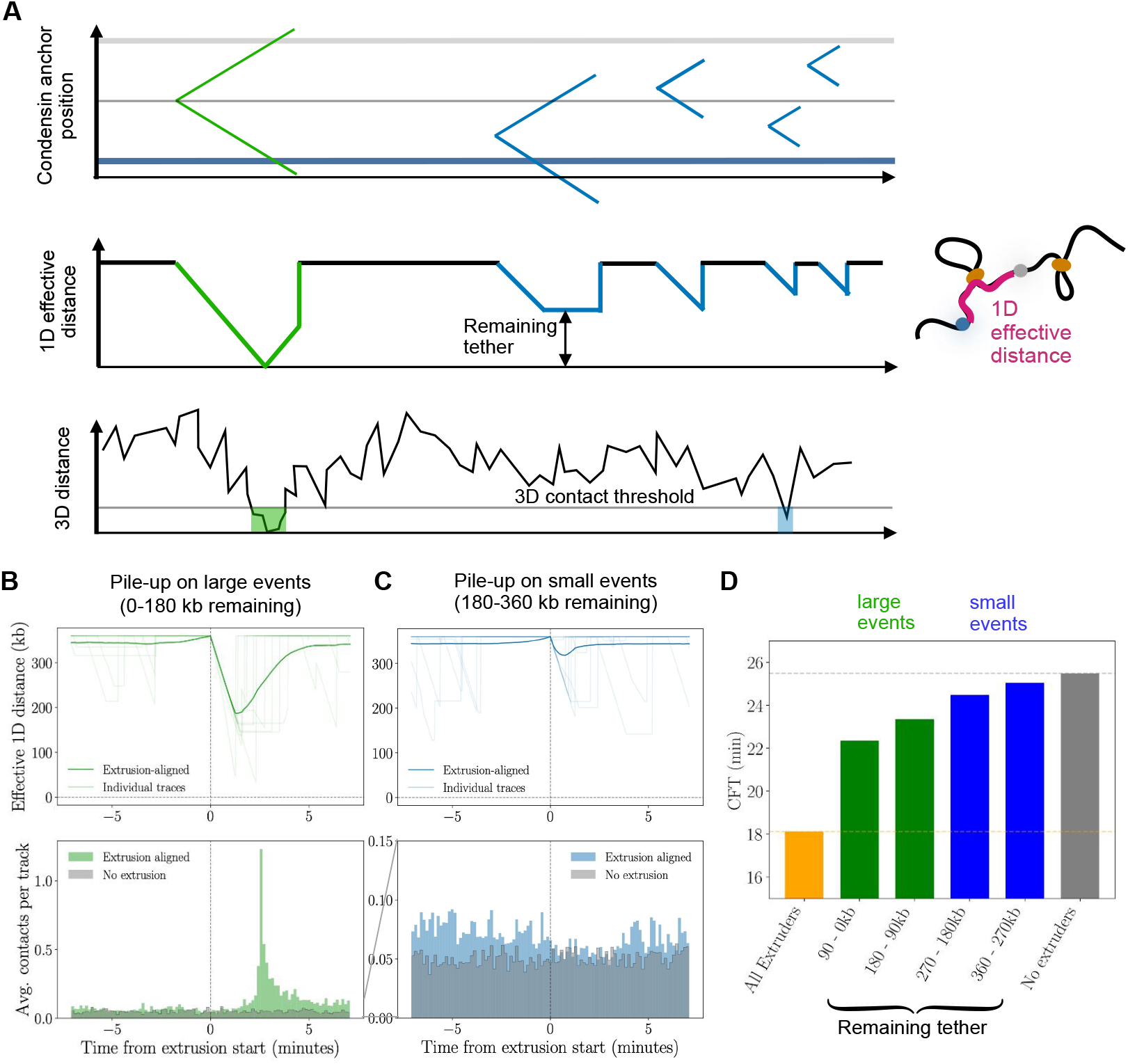
Two classes of extrusion-mediated processes accelerate long-range chromosomal contact formation. **A)** Schematic representation of both extrusion-mediated acceleration processes (green: large events, blue: small events). *Top*: Time series of the genomic region around the arrays. Each pair of lines represents the positions of the anchors of a single extruding condensin. *Middle:* Corresponding effective 1D distance between the arrays (defined schematically), showing shortening due to condensins landing in between. *Bottom*: Corresponding 3D distance between arrays with 3D contact highlighted. **B)** Top: Average effective 1D distance between the arrays aligned to the onset of large extrusion events (defined as remaining tether <180 kb). Faded lines show individual traces. *Bottom*: Average contact probability per track aligned to large extrusion event onset (green) compared to random times in simulations without extrusion (grey). High contact probability is present at 3 minutes (time to extrude to the arrays) following condensin binding. **C)** Same as B, but for small extrusion events (remaining tether > 180 kb). **D)** CFT simulation with specific extrusion events removed to quantify the contributions of large and small loops. For example, “0-90 kb remaining tether” reprents removing all extrusion events with remaing tether > 90 kb.

## Discussion

Functional long-distance chromosomal interactions in the native genome, such as looping between distal enhancers and their target promoters, tend to be rare and dynamic^37-39^. This dynamic nature coupled with limited time and spatial resolution of imaging technologies makes it very challenging to accurately detect interaction events. Consistent with this, polymer simulations have demonstrated that simple thresholding of the enhancer-promoter distance or Gaussian mixture modeling can lead to a substantial overestimation of the looped fraction^18^. In contrast, the interactions generated by CICI are mediated by dimerization between long arrays associated with hundreds of molecules. We expect such multivalent interactions to be stably maintained as long as the arrays are partially occupied. Indeed, while CICI junctions can be disrupted during anaphase due to chromosome segregation^13^, they remain remarkably stable in G1 cells, with virtually no disruption during our imaging period. This stability enables us to score interaction events and quantify first passage times with high confidence.

Our CFT measurements reveal that intra-chromosomal encounters in G1 yeast occur more rapidly than inter-chromosomal ones, and this difference relies on the presence of condensin, but not cohesin, consistent with their roles through the cell cycle of in *S*.*cerevisiae*. Depletion of condensin also selectively reduces long-distance contact frequencies within individual chromosomes. While we cannot directly observe condensin activity inside cells, we propose that loop extrusion is a highly plausible mechanism underlying the accelerated intra-chromosomal encounters due to the following reasons. First, condensin-mediated loop extrusion has been directly visualized *in vitro* using single-molecule assays^10^, indicating that condensin has loop-extrusion activities. Second, loop extrusion has the unique capability to differentially impact intra-vs inter-chromosomal interactions. Third, changes in the contact probability curves upon condensin depletion were consistent with the loss of extruded loops as seen in simulations. The selective effect on intra-chromosomal encounters would be difficult to explain through alternative mechanisms such as chromosome tethering through protein-protein interactions. Finally, incorporating the measured extrusion rates into our polymer model yields simulations that closely match both our CFT and Hi-C data, and provide estimates of condensin extrusion characteristics (e.g. velocity ∼2 kb/s), consistent with independent measurements of 1-3 kb/s^10,36^.

While previous studies have shown that condensin in budding yeast plays specialized roles at particular genomic regions, such as rDNA, the mating-type locus, and centromeres^20,40^, our analyses extend these observations by revealing condensin’s broader influence on 3D genome organization along regular chromosome arms. Our best-fit parameters of *d* and *λ* indicate that loops are relatively sparse in individual cells, with only ∼10% of the genome are extruded into loops at any given moment. The scarcity of chromosome loops is consistent with small changes in Hi-C signals. The condensin binding rate per unit genomic length can be estimated as *ν/λd*, corresponding to one condensin loading event every 1 to 10 minutes per Mb. Interestingly, even such low density of loop extrusion events, when combined with long processivity and high extrusion speed, is sufficient to promote long-range genomic communications. We find that this acceleration is primarily driven by rare, large extrusion events that leave the remaining separation between two loci below ∼100 kb. This mechanism likely applies to how cohesin-mediated loop extrusion facilitates enhancer-promoter communication in metazoans.

The observation that condensin, rather than cohesin, mediates loop extrusion in G1 yeast is in striking contrast with the well-established role of these two complexes in metazoans. In these higher eukaryotes, cohesin is the primary loop extruder during interphase, and condensin is mostly responsible for compacting chromosomes during mitosis. Notably, this canonical division of labor does not even apply to budding yeast during mitosis, where cohesin, instead of condensin, drives the genome-wide compaction of mitotic chromosomes^27,31^. The reason behind this functional divergence is unknown.

Nevertheless, cohesin loop extrusion in higher eukaryotes may also lead to selective acceleration of intra-chromosomal encounters. Many well-characterized distal enhancers and their target genes—such as the locus control region and the β-globin genes—reside on the same chromosome^41^. Similarly, long-range promoter-promoter interactions in *Drosophila* tend to occur within individual chromosomes^42^. Our findings suggest that loop extrusion may regulate gene expression by modulating the frequency of encounters between genes and their distal regulatory elements. Supporting this idea, cohesin activity has been shown to vary in neurons to control the expression of cell surface proteins by altering their enhancer-promoter interactions^43^. Moreover, our results in **Figure 2F** indicate that the enhancement of encounter rates by loop extrusion is distance-dependent. Intra- and inter-chromosomal encounter rates converge for pairs separated by small 3D distances, suggesting that interactions between proximal loci occur predominantly through diffusion. An active, ATP-driven process like loop extrusion is only required to promote encounter between distal loci. Consistently, cohesin was found to be dispensable for genes activated by nearby enhancers, but is essential for the full expression of genes regulated by distal enhancers^44,45^.

Unlike higher eukaryotes, budding yeast genes generally lack distal enhancers, yet a few hundred genes exhibit mild expression changes upon condensin depletion in G1^46^. These genes tend to be located near condensin-binding sites^46^, suggesting that condensin depletion may disrupt the 3D organization of these genes, e.g. by altering their contacts with other loci. Given that gene clustering in yeast has a modest effect on gene expression^47,48^, it is plausible that some of the gene expression changes result from altered 3D genome conformation. The detailed mechanism underlying these effects requires further investigation.

Loop extrusion by condensin may have functions beyond gene regulation, and the best example is its well-documented function in regulating mating type switch in G1 yeast. This is further supported by our Hi-C data showing significantly reduced contact frequencies along ChrIII in the absence of condensin. Together with impaired mating type switch efficiency in condensin-depleted strains^24,29,30^, this supports a model where condensin accelerates and stabilizes the interactions between the *HML* and the *MAT* locus and therefore promoting their recombination. A similar principle applies to V(D)J recombination during mouse B cell development, where cohesin-based loop extrusion is modulated to generate different contacts between the VH genes and the recombined DJH segments to diversify antibody repertoire^49^.

Another potential function of condensin-mediated loop extrusion is to promote disentanglement and individualization of chromosomes. This is evidenced by our Hi-C data near centromeres and rDNA, regions where condensin is highly enriched. In the absence of condensin, these regions exhibit higher inter-chromosomal contacts, indicating greater chromosomal intermingling. Correspondingly, our CFT data reveal that, while condensin depletion primarily slows intra-chromosomal encounters, it also mildly accelerate some inter-chromosomal interactions. Together, these results suggest that condensin promotes spatial separation of chromosomes, which may contribute to their proper segregation during mitosis. Consistent with this, in higher eukaryotes, condensin depletion leads to reduced territorial organization of interphase chromosomes and impaired separation of mitotic chromosomes^35,50-52^.

## Methods

### Plasmid and strain construction

We used the same background strain and the plasmids as described in our previous works^13^. Specifically, in this study, we started with the background strain containing *tor1* and *fpr* mutations, *REV1pr-LacI-GFP* and *REV1pr-TetR-mCherry* were inserted into *HIS3* locus, and *REV1pr-LacI-FKBP12* and *REV1pr-TetR-FRB* were inserted into the *ADE2* locus. For the two arrays, we started with the plasmids containing 192X TetO repeats (pSR11) and 256X LacO repeats (pSR13) as the template. We assembled homologous sequences from various loci on ChrIV and ChrXV with the two template plasmids. The reconstructed plasmids were cloned in stable *E. coli* (Invitrogen MAX Efficiency Stbl2 Competent Cells, 10268019) and were checked for length to be the same as original template. The plasmids were then transformed into yeast background strain. After every step of transformation, the insertion loci were confirmed by PCR. Each strain was checked under fluorescence microscope to confirm that it contains only one pair of red and green dot in G1. For strains with auxin-induced degradation, we tagged the C-terminus of endogenous Smc1 or Smc4 with AID and a V5 tag, and introduced OsTIR1 to the cells^53^. Depletion of the tagged protein was achieved by adding IAA (Sigma, I2886) to a final concentration of 500 μM. Strains were incubated with IAA for 1 hr prior to microscopy measurement. Depletion was confirmed by western blot. See **Table S1** for detailed plasmid and strain information.

### Fluorescence microscopy

CICI strains were cultured in synthetic media SCD to log phase, lightly sonicated to disperse cell clusters, and then incubated in SCD + 5 μM alpha factor (Zymo, Y1001) for ∼2.5 hrs to arrest the cells in G1 phase. For experiments involving auxin-induced degradation, cells were first treated with α-factor for 1.5 hrs, followed by the addition of IAA for another 1 hr. G1 arrest was confirmed microscopically by the presence of shmoo formation in a large fraction of cells. G1-arrested cells were then placed onto 1.2% agarose pads prepared with or without 1 ng/μl rapamycin (Sigma, 53123-88-9). We imaged these samples using our published time-lapse microscopy protocol^54^. For the + rapamycin samples, we typically started imaging within 5-10 min of rapamycin application. For each frame, three images were acquired sequentially in the order of phase, mCherry, and GFP fluorescence.

For the –rapamycin condition, two imaging acquisition settings were used. In the faster acquisition mode, two z-stacks were captured with 0.7 μm spacing and 0.1 s fluorescence exposure. Each fluorescence channel required ∼5 s to finish, and the time interval between frames was 10 seconds. In the slower acquisition mode, the same spacing and exposure were used, but with three z-stacks, and the interval between frames was extended to 120 s. For the +rapamycin condition, the time of initial rapamycin exposure was defined as t = 0 min. Imaging was performed using seven z-stacks with 0.6 μm spacing and an exposure time of 0.12–0.15 s. Each fluorescence channel took approximately 18 s, and the interval between frames was set to 240 s.

### Image analysis

We used the same MATLAB software developed in our previous works to annotate cells, detect dot positions and dot distances^47,54^. Briefly, cell contours were identified based on phase contrast images. For each pixel within a cell boundary, the program scanned across the z-stacks and recorded the maximum fluorescence intensity for both mCherry and GFP channels. These maximum intensity values were compiled into a 2D projection image for each channel. Fluorescent dots were then identified as pixel clusters with intensities significantly above background, and their coordinates were recorded. All detected dots were manually reviewed, and visually incorrect assignments were removed, especially when more than one red or green dot per cell were detected. The distances between mCherry and GFP dots in each cell were then calculated as distances on a 2D plain.

To identify CFT, we applied the following procedure. First, we excluded traces that met any of these criteria: interrupted in more than 60% of frames, lacked detectable signal for over 1 hr, or contained fewer than six valid data points. Second, we scanned the remaining traces for continuous colocalization events, defined as dot distances below 0.38 μm for at least four consecutive frames. If such a pattern was detected, the first frame in which it occurred was designated as the CFT. We applied this method to -rapamycin condition and detected CFT in 0.9% of traces which was the false discovery rate. Third, we assessed whether disassociation occurred after CFT by identifying the first frame in which the dot distance exceeded 0.5 μm following initial colocalization. In the majority of cases, colocalization persisted after CFT. In traces showing multiple rounds of association and disassociation, only the initial association event was counted to avoid double-counting.

### RMSD before and after CICI formation

To quantify the diffusion properties of the chromosome arrays, we used relative mean squared displacement (RMSD), which is defined as

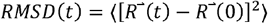

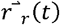 and 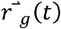 are the position of red and green dot respectively. 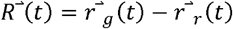 is the relative displacement between the two dots. If we ignore the correlations between 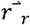 and 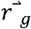,

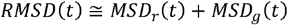

The RMSDs were derived from time-lapse imaging with three different frame rates. We averaged over all location datapoints separated by the same t. The fitting of RMSD before CICI formation agrees well with the Rouse model, where *MSD*(*t*) = 6*k*_*B*_*T*(*πKγ*)^-1/2^*t*^1/2^.

After CICI formation, the two arrays ideally should be at the same position: 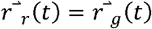 and thus 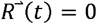. However, in our time-lapse imaging, the mCherry and GFP fluorescence channels were not imaged simultaneously, and therefore 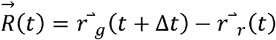. This time lag Δt depends on the number of z stack in our time-lapse imaging, and for +rapamycin condition, it is about 18 s. Substituting 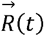 into the RMSD equation, we get:

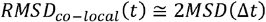

In other words, RMSD after CICI formation is a constant determined by the total diffusions of two arrays at time Δt, which can be compared with RMSD before CICI formation at t= Δt (**Figure 1G**). In Rouse model, when two beads come together, the friction coefficient of the dimer y is twice the monomer. The spring constant K is also doubled. Therefore, we expect the coefficient of MSD in Rouse model, 6*k*_*B*_*T*(*πKγ*)^-1/2^, to decrease by two folds. That is indeed what we observed experimentally.

### Prediction of the first passage time

MSD of Rouse motion is *Dt*^1/2^, where *D* is the diffusion constant. This means that the array explores roughly a spherical region with radius of *D*^1/2^*t*^1/4^ within time *t*, which consists of lattice sites ∝ *D*^3/2^*t*^3/4^. Meanwhile, the lattice sites visited by the dot should be ∝ *t*. In long time limit, the array visits each site infinitely often, which is known as “compact exploration”^55^. Therefore, the array should interact with its target once its explored radius is comparable to the distance between them: *d* = *D*^1/2^*t*^1/4^. In other words, CFT should have roughly a quartic dependence on the distance.

### Hi-C experiments

Hi-C was adapted from a previously published protocol^30^. Yeast were incubated in 200□mL SCD medium until OD660□reached□0.3. Cells were fixed with 3% formaldehyde (Fisher, RSOF0010-250A) for 20□min at 25□°C and then quenched with 0.2 M glycine (Fisher, 194825) for 20□min at room temperature. Cells were collected by centrifugation and washed with SCD. Cell pellets were resuspended in 1□mL TBS (Tris-buffered saline), 1% Triton X-100 (Sigma, 9036-19-5) and 1X protease inhibitor cocktail (Thermo Fisher, 87786). Cell lysate was generated by adding 500□μL acid-washed glass beads (Sigma, G8772) and vortexing for 25□min at 4□°C. Chromatin was recovered through centrifugation, washed with 1□mL TBS, resuspended in 500□μL 10□mM Tris-HCl buffer and digested with DpnII (NEB, R0543L) overnight at 37□°C. Digested DNA fragments were filled in with biotin-labeled dATP by incubating with Klenow enzyme (NEB, M0212), biotin-14-dATP (Thermo Fisher, 19524016), dCTP, dTTP, and dGTP (Thermo Fisher, R0181) for 4□hrs at room temperature. The biotin-filled DNA fragments were ligated by T4 DNA ligase (NEB, M0202L) for 4□hrs at 16□°C. Crosslink was reversed by incubation with proteinase K (Thermo Fisher, EO0492) at 65□°C overnight. DNA was purified by phenol-chloroform extraction. Biotin-labeled, un-ligated fragment ends were removed by incubating with T4 DNA Polymerase (NEB, M0203), dATP and dGTP for 4□hrs at 20□°C. DNA was cleaned by DNA clean and concentrator-5 kit (Zymo, D4014) and sheared by Diagenode Biorupter Pico (EZ mode, 30□s on, 30□s off, 15 cycles). Biotin-labeled DNA was enriched by MyOne(tm) streptavidin C1 beads (Thermo Fisher, 65001). Hi-C libraries were prepared with NEBNext Ultra II DNA Library Prep Kit. Next generation sequencing was performed on NextSeq 2000, and 100□million of 50□bp paired-end reads were generated for each replicate. Hi-C data analysis was performed with HiC-pro (version 3.1.0)^56^, cooler (version 0.10.0)^57^, cooltools (version 0.7.0)^58^ and HiCExplorer (version 3.7.2)^59^.

### Loop extrusion simulation

#### Polymer simulation

Polymer simulations were conducted using OpenMM and the polychrom library (http://github.com/mirnylab/polychrom). Each simulation comprised ten freely-jointed polymers, each consisting of 1800 beads, where each bead represents 1 kb of chromatin. The polymers were confined within a spherical potential with a radius 50 times the bond length. Monomeric subunits were connected into a linear polymer through pairwise harmonic bond interactions. Condensin positions, simulated independently as described below, introduce additional harmonic bonds bridging specific monomers. Monomeric subunits repel each other through weak excluded-volume-like interactions. The viscosity of the implicit solvent was set to ensure Brownian dynamics across time scales longer than the shortest polymer relaxation time.

#### Condensin dynamics simulation

Simulations of condensin were performed using the lefs-cython library (https://github.com/mirnylab/lefs-cython/). Chromatin is modeled as a 1D lattice, with each site representing a monomer from the 3D simulation. Each chromatin-bound condensin occupies two lattice sites, corresponding to its anchor points on the DNA. The total number of condensins, *N*, present in each simulation is constant and determines the typical distance between two condensins (i.e. centers of the loops)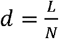, and the linear density of condensin is given by 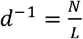. Bound condensins extrude symmetrically, at a constant velocity *ν*. Once extrusion has started, condensins unbind DNA at a rate *k*_*u*_. When the heads of two condensins collide, they block each other at the site of collision while the other head continues extruding. If a condensin anchor reaches one end of a polymer, it is unloaded. Each time a condensin unbinds, another one binds at a random location. Consequently, the binding rate per lattice site of condensin, *K*_*b*_, is related to the condensin density and condensin unbinding rate through the relation 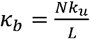. The processivity, i.e. the average loop length of unobstructed condensin,is given by 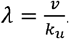.

#### Simulations with size-selective extruder removal

To isolate the effects of different extrusion event sizes, we performed simulations where extruders where selectively removed based on the remaining length of DNA tether between the arrays during the extrusion process. We defined an extrusion event as a contiguous moments in time where the effective 1D distance between the arrays drops below its equilibrium value due to one or several loop extruder. Using extrusion tracks from simulations that best fit the experimental data, we identified all such events and categorized them by the minimal remaining 1D distance between the arrays reached during the event (**Figure 7A**). We then removed all extruders involved in extrusion events outside the desired range and re-ran the polymer simulation with the filtered extrusion track.

#### Calibration of Time and Length Scales

Simulations were conducted without extruders, initially using arbitrary time and length scales. These were later rescaled against the extrusion-free (condensin-depleted) experiments. The rescaling was performed by matching the two-point MSD of two beads in the simulation to the experimentally measured two-locus MSD, with bead separation in the simulation chosen to correspond to the genomic separation of the pair in the experiment.

#### Simulation CFT and Array Reaction Radius Calibration

To reproduce CICI formation time quantitatively, we compute the average time for the distance between bead pairs to drop below a threshold value, representing the interaction radius of the arrays, starting from a randomly chosen time point that represents the time of rapamycin introduction. The reaction time depends on the reaction radius - a larger reaction radius makes it more likely for arrays to encounter and react, thus reducing the reaction time. Since the interaction radius of the arrays is unknown, we determine its value by computing the simulation CFT across a range of interaction radii and selecting the value that best matches the experimental CFT. We averaged the CFT across multiple pairs of monomers with the same bead separation, taking advantage of the statistical equivalence of simulated polymers and their translational symmetry along the polymer backbone. Again by symmetry, since our simulations neglect peripheral tether, all pairs of monomers located on different polymers are statistically equivalent (modulo boundary effects). The CFT for trans pairs was computed from randomly selected pairs of monomers located on different polymers.

#### Contact probability

Contact frequency curves were computed from the simulated conformations using the polykit library (**https://github.com/open2c/polykit**).

## Data availability

The Hi-C sequencing data generated in this study have been deposited into the Gene Expression Omnibus (GEO) database under accession code GSE296183.

## Acknowledgements

We acknowledge all members in Bai lab and Mirny Lab for insightful comments on the manuscript. We also want to thank the members of the Center of Eukaryotic Gene Regulation at PSU for discussions and technical support. This work is supported by the National Institutes of Health (T32 GM125592 to Y.L., R35 GM139654 to L.B., NIH R01GM114190 to L.M.) and the National Science Foundation (MCB-2016266 to L.B., NSF 2210558 to L.M., NSF 2044895 to L.M.). LM is also a Simons Investigator.

## Author Contribution

F.Z. and Y.L. designed and performed most of the experiments; C.S. contributed to the experiments; F.Z., Y.L. and L.B. performed data analysis and wrote the manuscript. T.F., H.D.P and L.M. performed and analyzed the polymer simulations.

---

**Figure S1.**
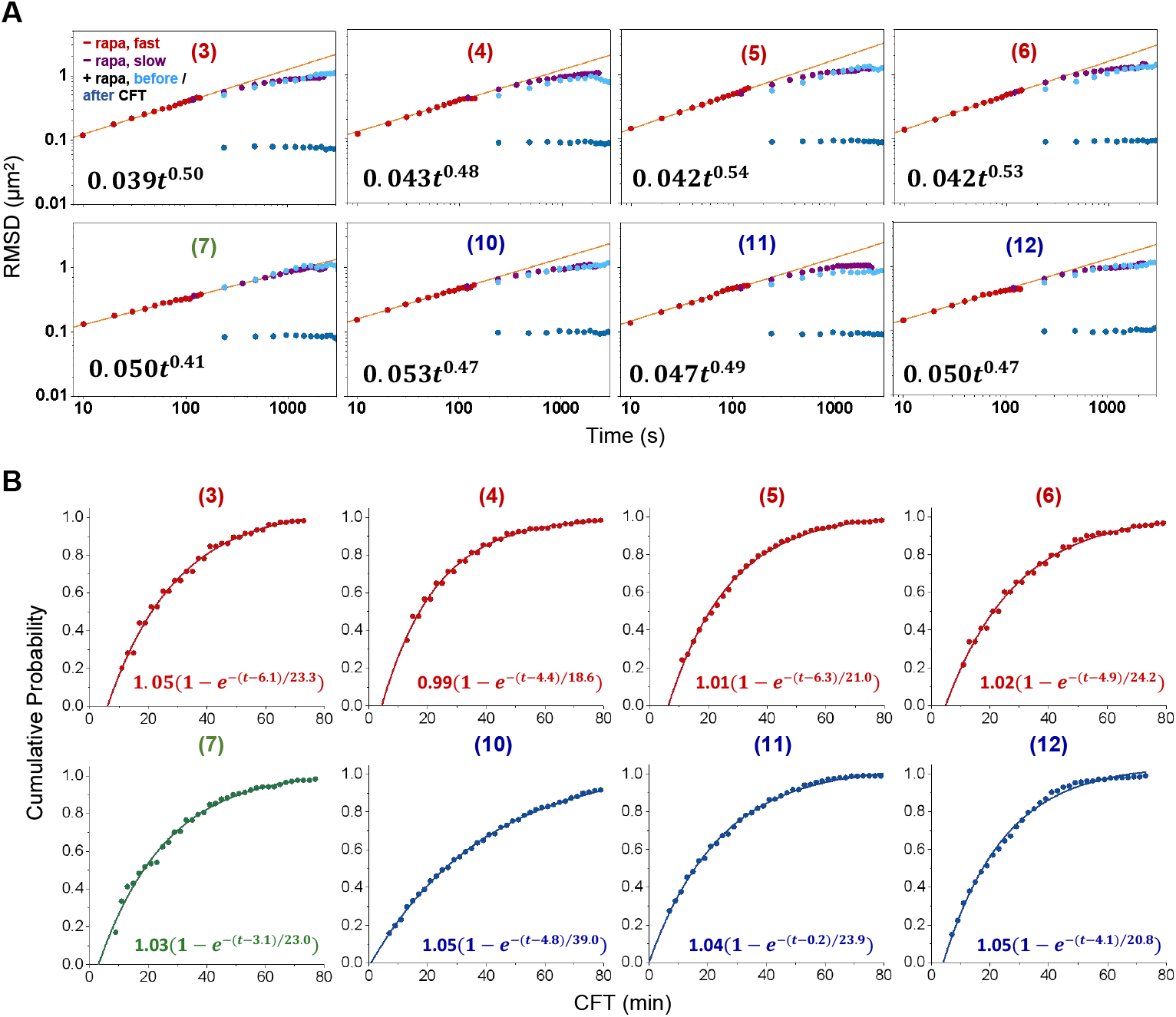
RMSDs and CFTs of more loci pairs. **A)** RMSDs as a function of time for intra-arm pairs 3-6, inter-arm pair 7, and inter-chromosomal pairs 10-12. Best power-law fits for the shorter time points are shown in the diagram. **B)** Cumulative CFT probabilities for the same pairs in A. The single exponential fit is shown in each panel.

**Figure S2.**
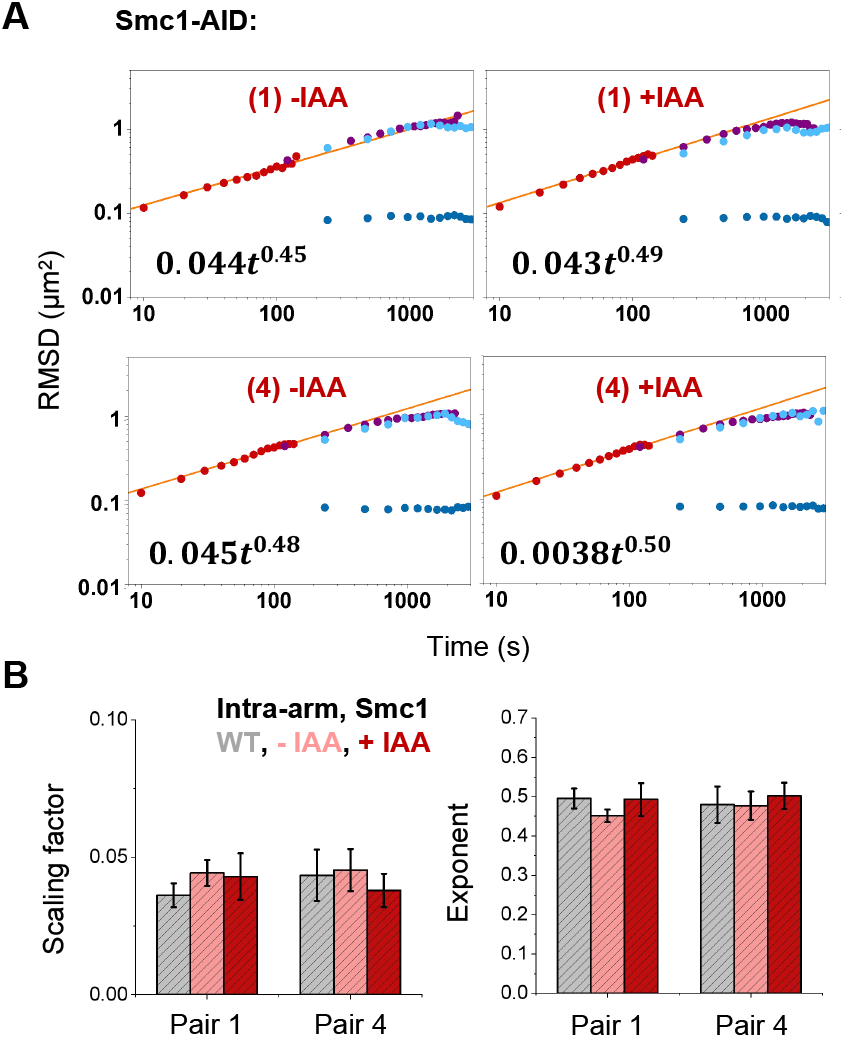
Smc1 depletion has little effect on chromosome motion. **A)** RMSDs as a function of time in Smc1-AID strain ±IAA for pair 1 and 4. **B)** The best-fit scaling factor and exponent of the RMSDs in WT and Smc1-AID strain ±IAA for pair 1 and 4.

**Figure S3.**
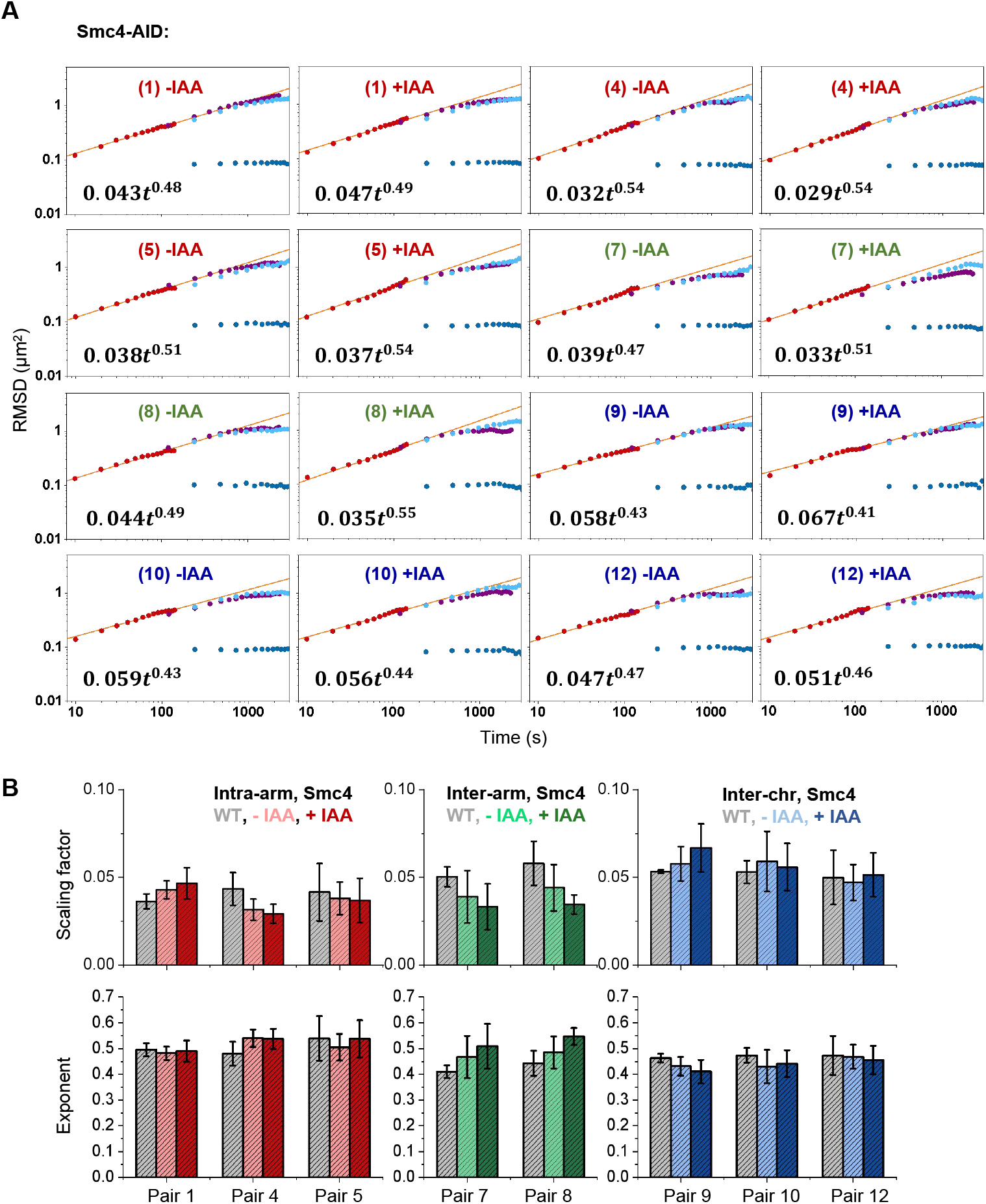
Smc4 depletion has little effect on chromosome motion. **A)** RMSDs as a function of time in Smc4-AID strain ±IAA for indicated pairs. **B)** The best-fit scaling factor (top) and exponent (bottom) of the RMSDs in WT and Smc4-AID strain ±IAA for indicated pairs.

**Figure S4.**
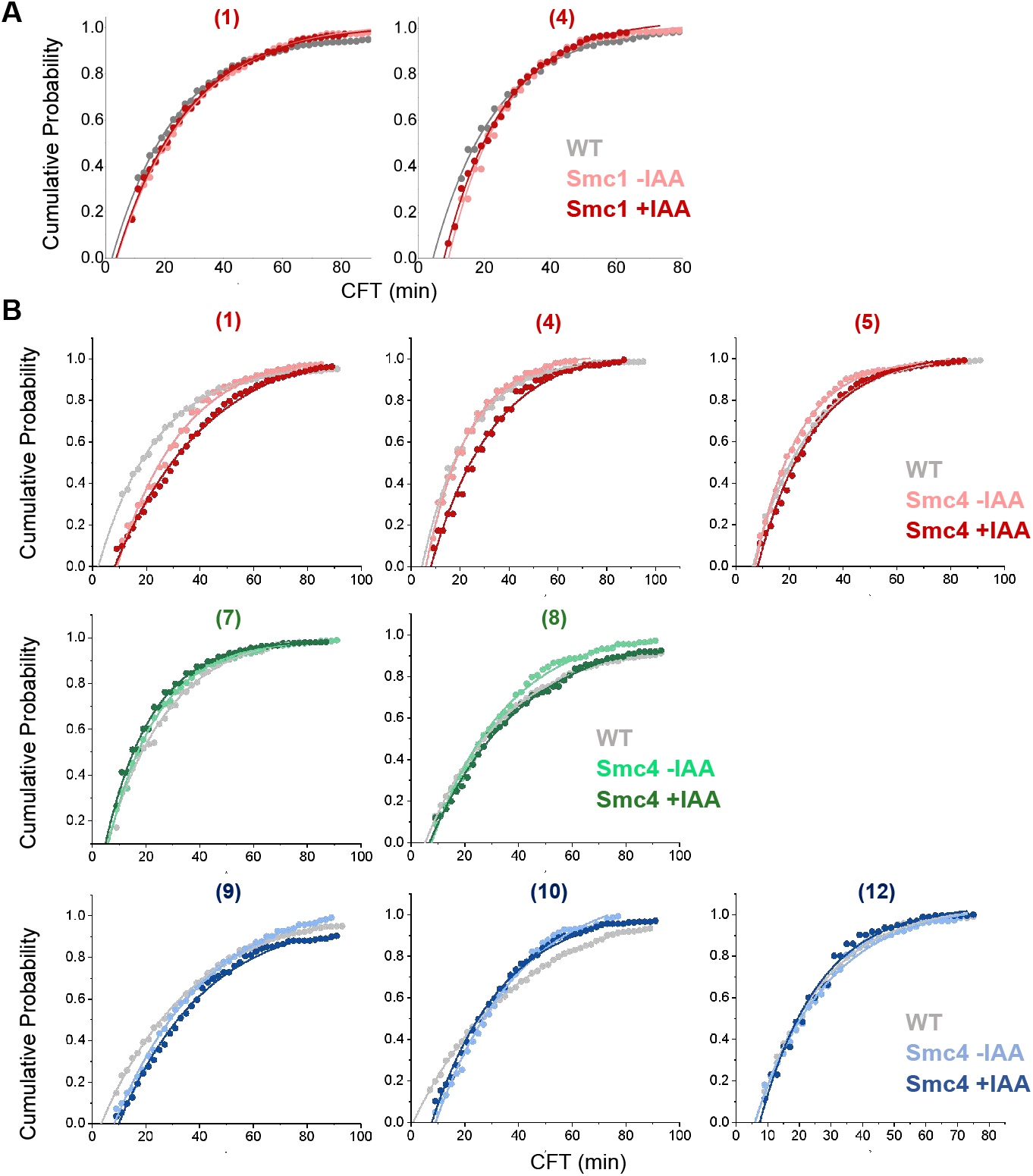
CFTs in Smc1/Smc4 depletion strains. **A)** Cumulative CFT probabilities in WT and Smc1-AID strain ±IAA for pair 1 and 4. **B)** Cumulative CFT probabilities in WT and Smc4-AID strain ±IAA for indicated pairs.

**Figure S5.**
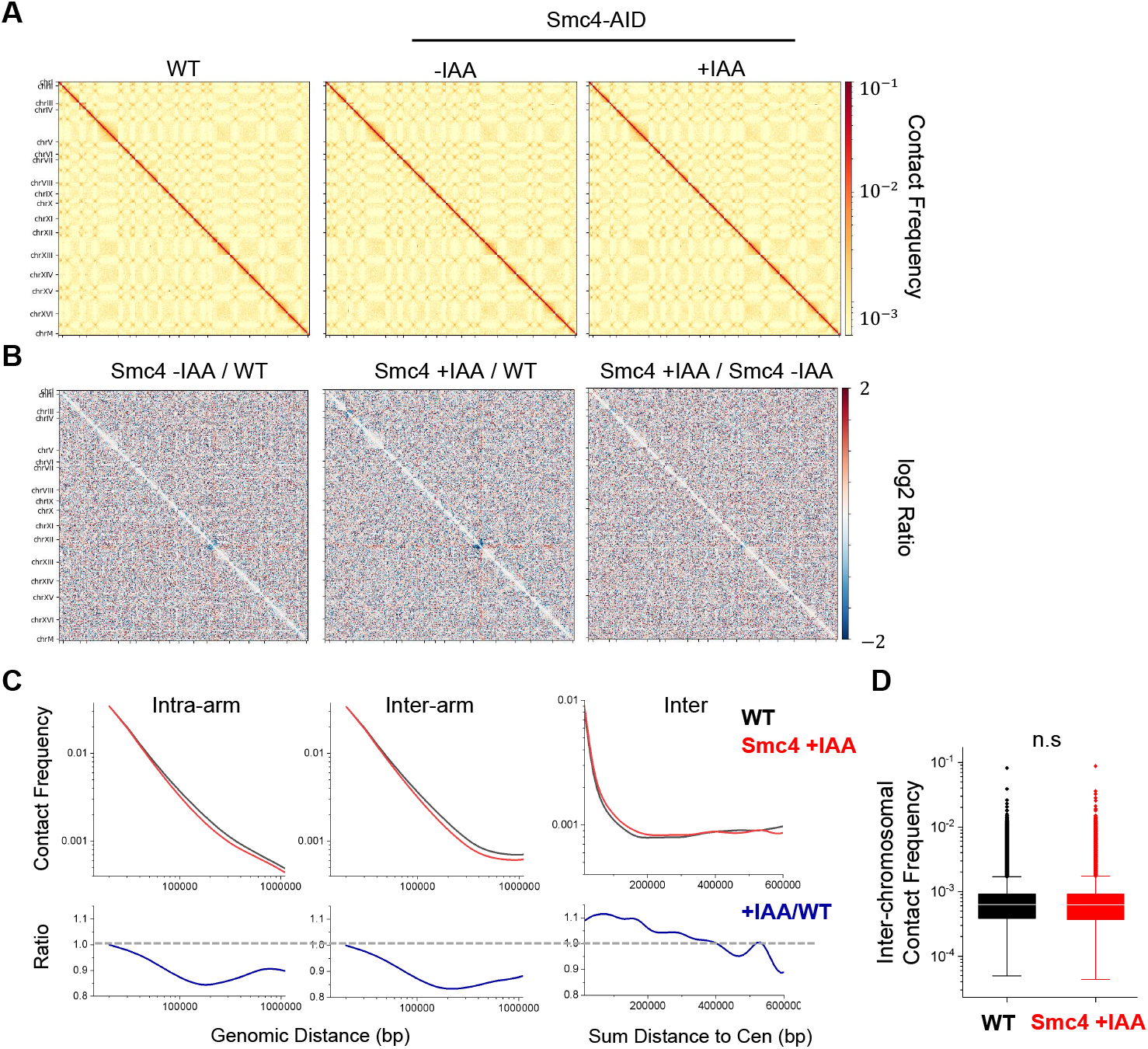
Condensin effect on genome-wide chromosome conformation in G1 yeast. **A)** Genome-wide Hi-C contact matrices in G1-arrested WT and Smc4-AID strain ±IAA. **B)** log2 fold change of the data in A. **C)** Contact frequency vs linear genomic distance in WT (black) and condensin-depleted (red) G1 cells. Upper panels show the contact frequencies averaged across the genome as a function of genomic distances, and the lower panels show the Smc4 +IAA / WT ratio. The left two columns represent loci pairs on the same chromosome arms or on the same chromosome but on different arms. The right column represents inter-chromosomal loci pairs with the same distance (x axis) to their corresponding centromeres. **D)** Contact frequencies between all possible inter-chromosomal loci pairs in WT and Smc4-depleted cells.

**Figure S6.**
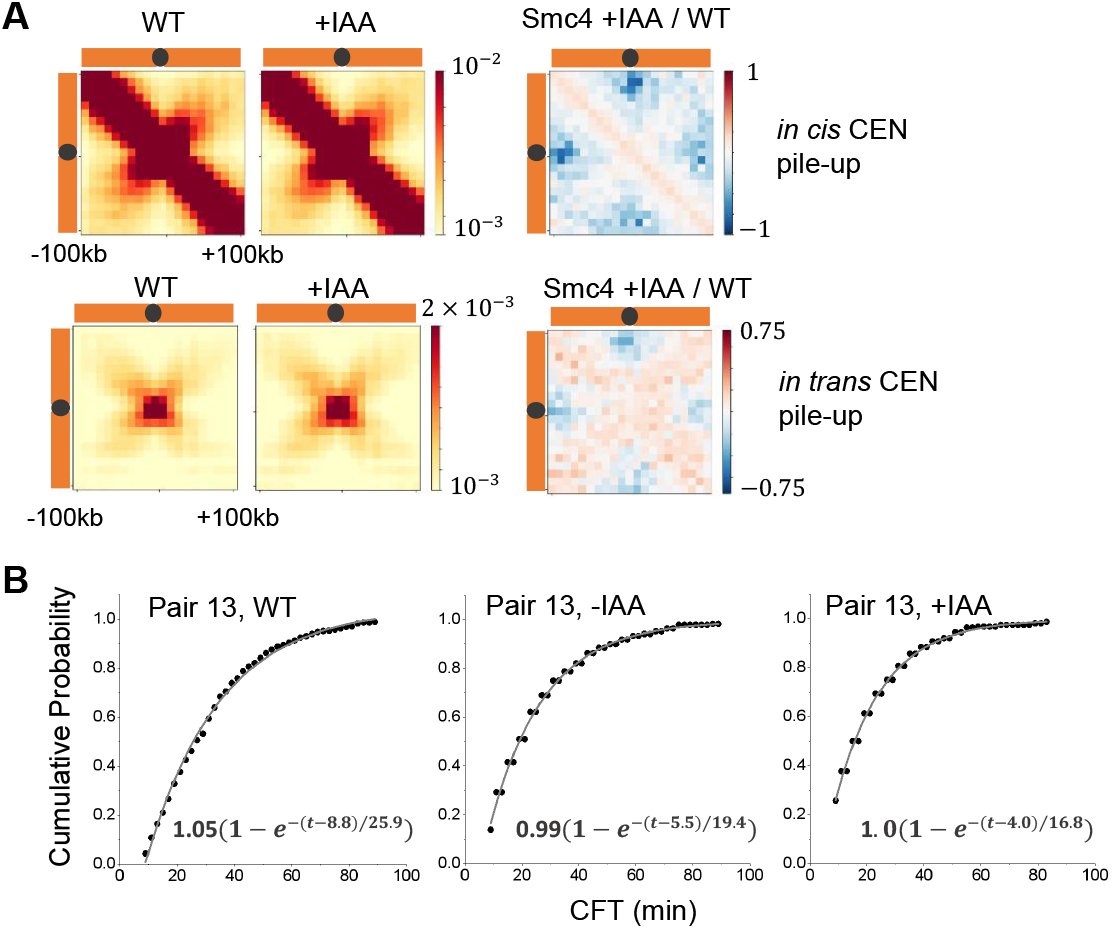
Condensin effect on Hi-C and CFT in specialized genomic region. **A)** Pile-ups of the Hi-C contact matrices of ±100kb regions centered around centromeres in in WT vs Smc4-depleted cells. Log2 fold change between Smc4-AID +IAA and WT is shown on the right. **B)** Cumulative CFT probabilities in WT strain and Smc4-AID strain ±IAA for pair 13.

**Figure S7.**
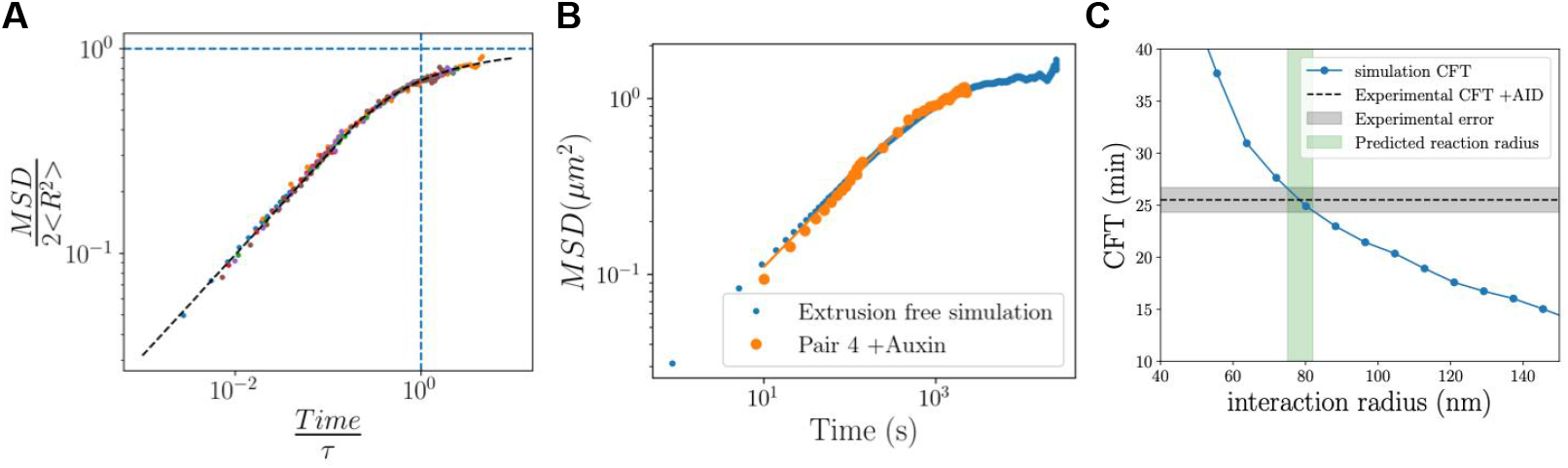
Rescaling procedure for mapping simulation length scales to experimental data. **A)** Experimental two-point MSD data for multiple WT and condensin depleted pairs, collapsed onto a single curve by rescaling time with the fitted Rouse time and MSD with the squared fitted spatial separation, *R*^2^ .The rescaling parameters, Rouse time and *R*, were obtained by fitting the theoretical two-point MSD for a Rouse chain (*M*_2_(*t*)) equation below). **B)** Experimental RMSD for locus pair 4 in condensin-depleted cells, overlaid with the corresponding rescaled simulation data. This comparison ensures that the simulation units are properly scaled to match the experimental RMSD. **C)** Search time computed in simulations without extrusion for different values of the array interaction radius. The gray dashed line represents the experimental search time for locus pair 4 in condensin-depleted cells, with the shaded region indicating the corresponding error bar. The vertical green shaded region marks the fitted interaction radius, determined as the value that best matches the experimental search time.

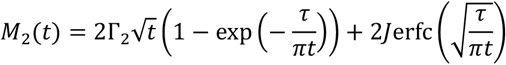

**Figure S8.**
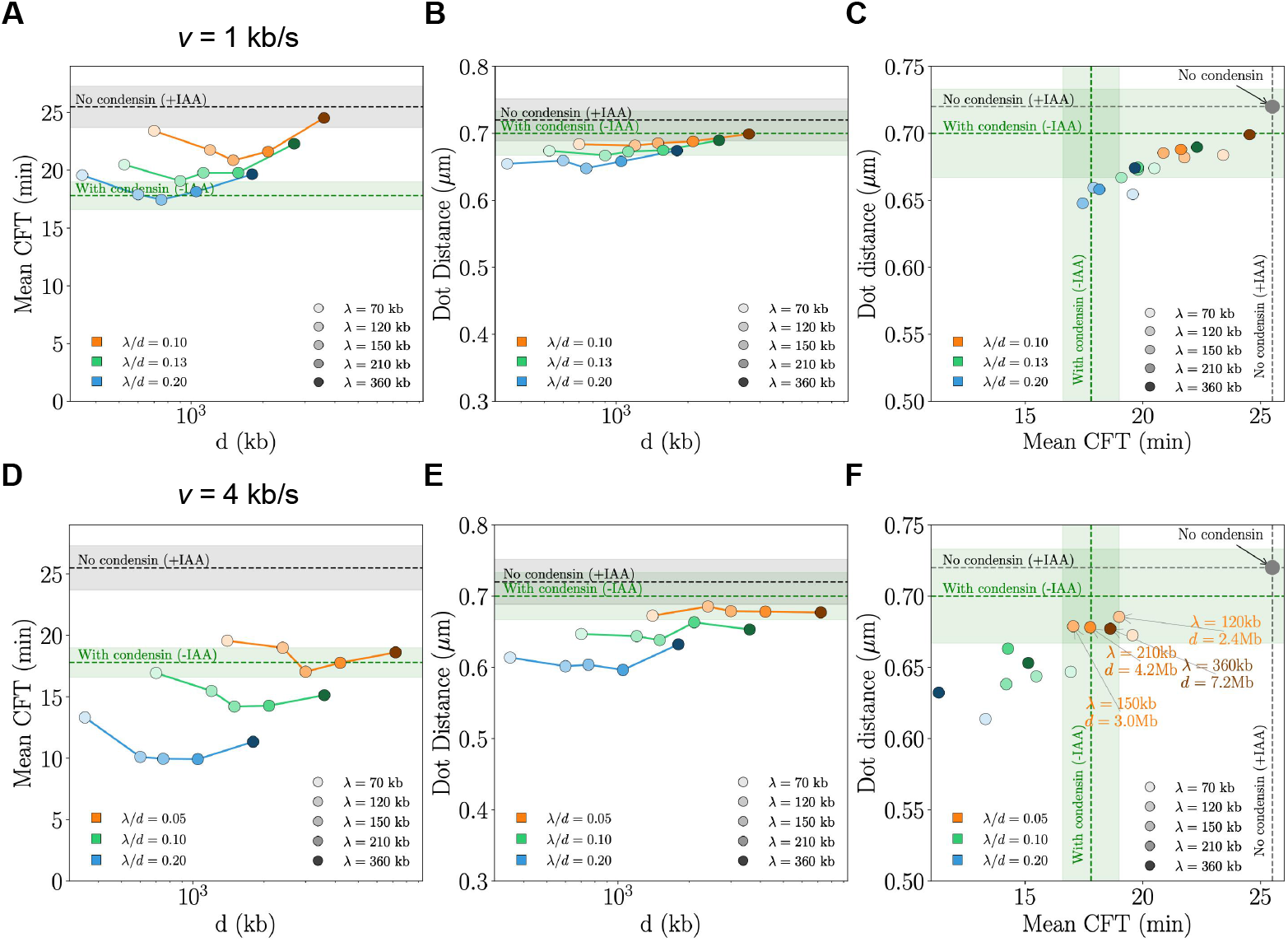
Simulated CFT and mean dot distance for extrusion speed of 1 and 4kb/s. **A)** The average search time between two monomers separated by 360 kb in simulated polymers with loop extrusion at *ν* = 1 *kb*/*s*, shown for different values of loop size (*λ*) and loop separation (*d*). Dashed lines indicate the experimentally measured search times in cells with condensin (-IAA, green) and without condensin (+IAA, grey), with shaded regions representing the corresponding error bars. **B)** Same as (A), but showing the average spatial distance between the two monomers instead of search time. **C)** The same simulation data points as in A) and B), but with dot distance plotted against search time. Simulations where both the dot distance and search time fall within 1.5 times the experimental error are annotated. **D-F)** Same than A-C, but for *ν* = 4*k*/*s*.

**Figure S9.**
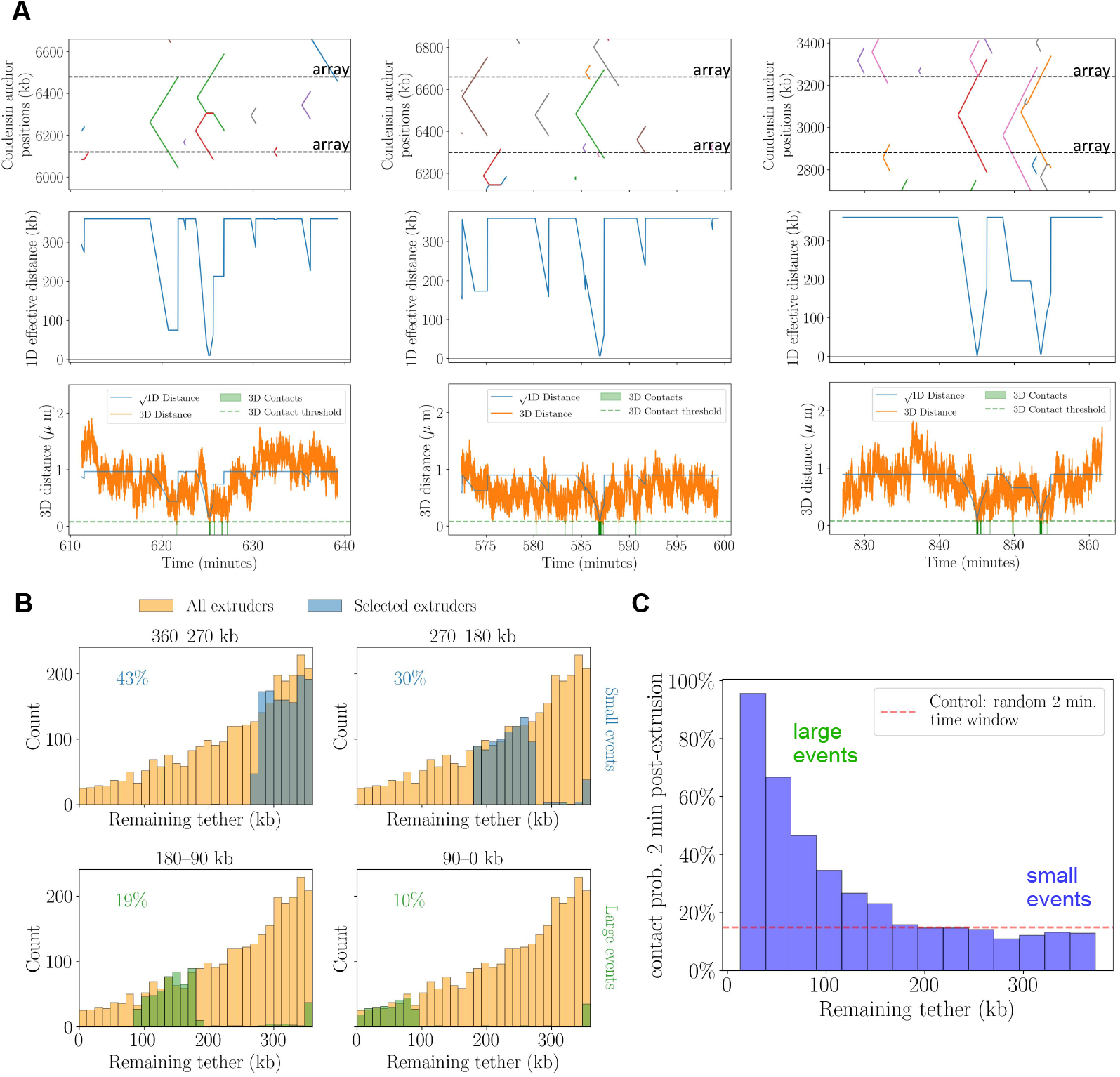
Contribution to CICI formation by large vs small extrusion events. **A)** Each panel shows representative simulated extrusion events where condensin reduces the effective 1D genomic distance between the arrays, triggering potential 3D contact. *Top*: positions anchored by condensin over time, with dashed lines marking the genomic positions of the arrays. *Middle*: 1D effective distance between the arrays. *Bottom*: corresponding 3D distance (orange), plotted alongside the re-scaled square root of the 1D distance (blue). Green shaded areas indicate periods where the 3D distance drops below the contact threshold. **B)** Histograms of minimal remaining tether during extrusion events. All extrusion events (orange) are shown alongside those retained for simulation under size-selective extruder removal (blue: small events; green: large events). Each panel corresponds to a different range polymer simulations isolating the effect of extrusion events of specific sizes (see Methods and Fig. 7A). **C)** Contact probability (in percents) within a 2-min window following the point of minimal 1D distance during extrusion events, binned by the minimal tether length remaining between the two loop anchors (x-axis). Blue bars represent the probability that the two loci come into 3D contact within a 2 minute time window after extrusion. Dashed red line indicates the baseline probability of contact at random times, computed by averaging contact formation over 2 minute random time windows.

